# FindZX: an automated pipeline for detecting and visualising sex chromosomes using whole-genome sequencing data

**DOI:** 10.1101/2021.10.18.464774

**Authors:** Hanna Sigeman, Bella Sinclair, Bengt Hansson

## Abstract

Sex chromosomes have evolved numerous times, as revealed by recent genomic studies. However, large gaps in our knowledge of sex chromosome diversity across the tree of life remain. Filling these gaps, through the study of novel species, is crucial for improved understanding of why and how sex chromosomes evolve. Characterization of sex chromosomes in already well-studied organisms is also important to avoid misinterpretations of population genomic patterns caused by undetected sex chromosome variation. Here we present findZX, an automated Snakemake-based computational pipeline for detecting and visualizing sex chromosomes through differences in genome coverage and heterozygosity between males and females. FindZX is user-friendly and scalable to suit different computational platforms and works with any number of male and female samples. An option to perform a genome coordinate lift-over to a reference genome of another species allows users to inspect sex-linked regions over larger contiguous chromosome regions, while also providing important between-species synteny information. To demonstrate its effectiveness, we applied findZX to publicly available genomic data from species belonging to widely different taxonomic groups (mammals, birds, reptiles, fish, and insects), with sex chromosome systems of different ages, sizes, and levels of differentiation. We also demonstrate that the lift-over method is robust over large phylogenetic distances (>80 million years of evolution).

## Background

Sex determination, the process by which sexually reproducing organisms initiate the developmental program to become male, female, or hermaphroditic, is remarkably diverse (Bachtrog 2014, Bull 1983, Beukeboom 2014). The mechanism that triggers the sex determination program can be either genetic, environmental, or a combination of both. In many animals and some plants this process is genetically controlled by sex chromosomes, where either males (XY systems) or females (ZW systems) are heterogametic, i.e., carry two different sex chromosomes. Sex chromosomes have evolved many times, both *de novo* in organisms without genetic sex determination and through turnovers of already existing sex chromosome systems (The Tree of Sex Consortium 2014; Bachtrog et al. 2014; Stöck et al. 2021). There have also been numerous translocations of autosomes to existing sex chromosomes (forming “neo-sex chromosomes”), which may result in variable gene content between the sex chromosomes of different, closely related species. Despite recent efforts in identifying and characterizing sex chromosomes across a broad range of taxonomic groups (e.g., Pease & Hahn 2012; Gamble et al. 2015; Jeffries et al. 2018), we are likely still missing much of the existing sex chromosome diversity. Filling these knowledge gaps is important for an improved understanding of how and why sex chromosomes evolve, and to avoid misinterpretations of population genomic data caused by undetected sex chromosome diversity.

Sex chromosomes can be identified using cytogenetic methods, where sex-specific karyotype differences (in chromosome number and/or size) can reveal the sex chromosome pair. These methods may, however, fail to reveal homomorphic sex chromosomes (as they are of similar size) and provide imprecise information on gene content and homologies to other species. Sex chromosomes can also be identified by contrasting genomic (or transcriptomic) data from males and females (e.g., Vicoso et al. 2013). This is because X and Y (or Z and W) are expected to evolve genetic differences as a consequence of recombination suppression. Such “sex-linked” regions may be detected through a range of genome signatures, including sex differences in allele segregation patterns, gene expression, heterozygosity, or genome coverage (reviewed in Palmer et al. 2019). Different data types, sampling strategies and computational methods may be suitable for detecting sex chromosomes of variable degrees of differentiation, with homomorphic sex chromosomes requiring more carefully designed computational methods and sampling strategies.

Several programs and computational resources for detecting sex-linked genomic regions have been published. Among these are SEX-DETector (Muyle et al. 2016; which uses expression data from a pedigree of samples), discoverY (Rangavittal et al. 2019; which finds sex-limited scaffolds through comparisons of reference genomes from a homogametic and heterogametic sample) and RADSex (Feron et al. 2020; which uses restriction site–associated DNA sequencing (RAD-seq) data from several males and females). Whole genome sequencing (WGS) is now a standard form of sequencing, and this kind of data is increasingly used to characterize sex chromosomes (e.g., Chen et al. 2014; Wright et al. 2017; Rafati et al. 2020). However, to our knowledge there is no published computational resource dedicated to detecting sex chromosomes using WGS data from males and females.

Here we present findZX, a Snakemake-based (Köster & Rahmann 2012) pipeline that identifies sex chromosomes, or more specifically the non-recombining part of the sex chromosomes, using WGS paired-end data from a flexible number of male and female samples. In essence, findZX scans across windows of a reference genome for two genomic signatures of sex-linkage: sex differences in (i) genome coverage and (ii) heterozygosity. The combined analysis of these measurements is a powerful and widely used approach for identifying sex chromosome systems of varying degrees of differentiation (Vicoso et al. 2013; Palmer et al. 2019). The pipeline can be applied to fragmented (scaffold-based) genome assemblies but also includes the option to perform a “lift-over” to a more contiguous reference genome of another species. The benefit of this is twofold: it allows users to inspect sex-linked regions over larger contiguous chromosome regions, while also providing detailed synteny information between species. FindZX is available on GitHub (https://github.com/hsigeman/findZX).

FindZX aligns paired-end WGS reads of female and male samples to a reference genome constructed from the homogametic sex (female XX or male ZZ) and detects chromosomal regions with sex differences in genome coverage and heterozygosity by contrasting them to the genome-wide (autosomal) pattern characterised by no such sex differences (Figure 1a). The sex-specific signature depends on the level of sex chromosome differentiation and Y/W degeneration because reads from the sex-limited chromosome (Y/W) may or may not successfully align to the homogametic (XX/ZZ) reference genome. In general, species with little differentiation between the sex chromosome copies (Figure 1b) are expected to show weaker sex-specific signals than species with highly differentiated sex chromosomes (Figure 1c,d). Restricting the number of allowed mismatches between the reference genome and aligned reads when attempting to identify sex chromosomes is recommended, as this prevents reads from the sex-limited chromosome aligning to its gametologous chromosome copy and thereby increasing the sex difference in genome coverage. For low differentiation (homomorphic) sex chromosome systems, the clearest genomic signature will be heterozygosity as many reads from the sex-limited chromosome (Y/W) will align (to the X/Z), while genome coverage may only reveal sex differences when strongly restricting the number of allowed mismatches (Figure 1b). For high differentiation (heteromorphic) sex chromosome systems, the clearest genomic signature will be coverage differences, especially when restricting the number of allowed mismatches (Figure 1c,d), while heterozygosity may sometimes be skewed towards the heterogametic sex if reads from the sex-limited chromosomes do align (Figure 1c) or towards the homogametic sex if they do not (due to genetic variation on X/Z resulting in heterozygosity only in homogametic individuals) (Figure 1d). Due to the use of a homogametic reference genome, findZX is designed to identify X and Z chromosomes/scaffolds rather than Y and W chromosomes. However, the output will reveal useful information about the size of the Y or W chromosome, and the level of X-to-Y, or Z-to-W, divergence.

**Figure 1.**
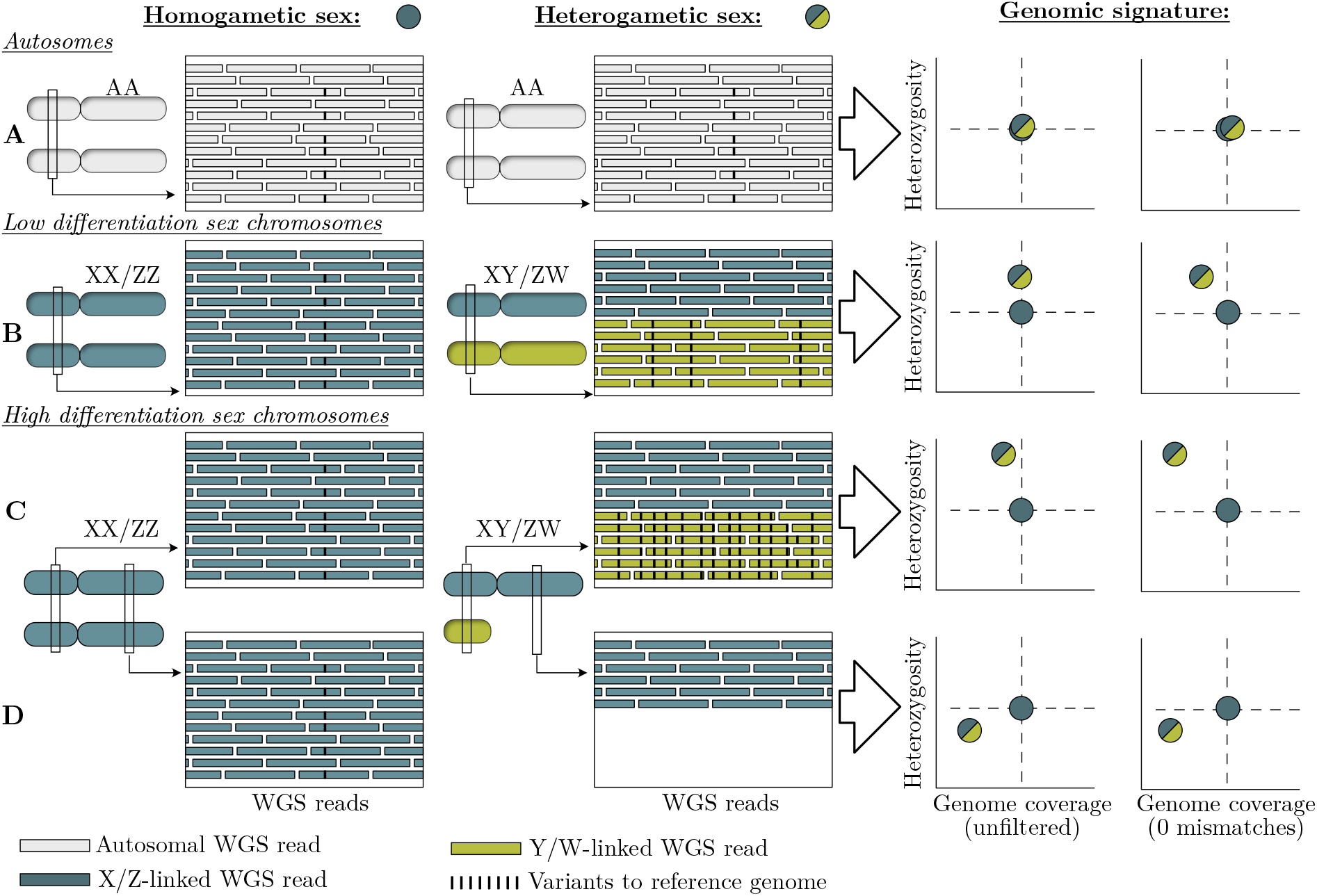
Sex-linked genomic regions of varying stages of differentiation can be detected through differences in genome coverage and heterozygosity between WGS reads of males and females aligned to a homogametic (XX/ZZ) reference genome. Reads are coloured according to chromosome origin (“autosomal”, “X/Z-linked” or “Y/W-linked”). Genome coverage is calculated as the number of WGS reads aligning to a specific genomic region. The black bars within the WGS reads represent genetic variants compared to the reference allele (i.e., heterozygous sites). (**A**) Genome coverage and heterozygosity is expected to be similar between sexes on autosomes. (**B**) Sex chromosomes of low differentiation are characterized by higher heterozygosity in the heterogametic sex. They often display equal genome coverage between sexes when allowing mismatching reads to map to the reference genome but pronounced coverage differences when restricting the number of allowed mismatches. (**C,D**) Highly differentiated sex chromosomes are expected to have either equal genome coverage between sexes when allowing mismatches, or lower genome coverage in the heterogametic sex if the sequence divergence is large enough to prevent successful alignment of reads to the reference genome. When restricting the allowed number of mismatches, we expect significantly lower genome coverage in the heterogametic sex. If the genomic region has a completely degenerated sex-limited chromosome copy, we expect lower genome coverage in the heterogametic sex regardless of the number of allowed mismatches. Highly differentiated sex chromosomes will have either (C) very high or (D) lower heterozygosity in the heterogametic sex depending on mapping success, level of Y/W degeneration, and level of genetic variation on X/Z.

We applied the pipeline to published data from species of various taxonomic clades and sex chromosome systems to demonstrate its effectiveness in both identifying the sex chromosomes and finding homology to other species.

## Results

### Pipeline overview – Input, computational steps, and output

FindZX identifies sex-linked genomic regions using WGS reads from samples of different sexes, with a minimum input of one individual per sex and a reference genome constructed from a homogametic individual (“study-species reference genome”). When no reference genome is available for the studied species (as would be the case for most studies), a scaffold-based assembly sufficient for the pipeline can easily be constructed *de novo* using the short-read data from one of the homogametic samples and standard assembly programs (see example on how to do this on the findZX GitHub page). Main computational steps (Figure 2a; see Methods for more details) and output plot types (Figure 2b-f) of the pipeline are shown in Figure 2. The pipeline can be run using either of two snakefiles: findZX or findZX-synteny. Input data and computational steps in blue boxes (Figure 2a; Steps 1-10) are common for both run modes. Computational steps in green are specific for findZX (Steps 11-12), whereas input data (a reference genome from a second species; “synteny-species reference genome”) and computational steps in orange (Steps 13-17) are specific for findZX-synteny.

**Figure 2.**
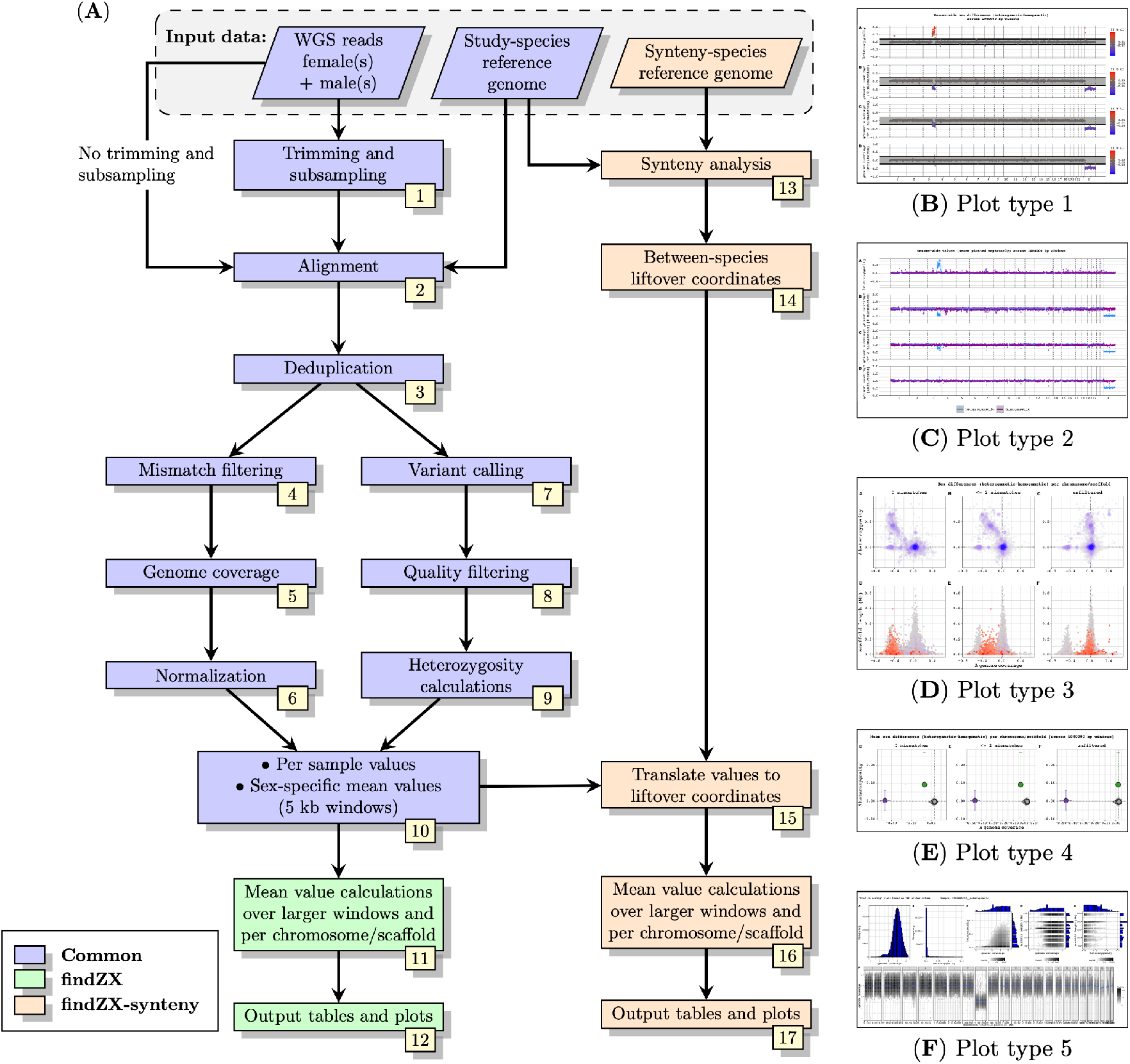
Flowchart of the main computational steps in the findZX/findZX-synteny pipeline (A), and miniatures of the five output plot types (B-F). **A)** The flowchart boxes in blue are common for both findZX and findZX-synteny. The green boxes are specific for findZX and orange boxes are specific for findZX-synteny. Parallelograms (top row) represent input data, and rectangles represent computational steps and output. See Main text and Methods section for details. Plot type 1 (**B**; see also Figure 3) and 2 (**C**; see also Supplementary Figure 2) show genome-wide heterozygosity and genome coverage values based on means across genome windows (here 1 Mb). Plot type 1 (B) shows sex differences and 2 (C) shows values for each sex separately. Plot type 3 (**D**; see also Figure 4) shows heterozygosity and genome coverage values for each chromosome/scaffold, as well as chromosome/scaffold length. Plot type 4 (**E**; see also Figure 5) shows mean ± SD sex differences (calculated from genome windows, here 1 Mb) for each chromosome/scaffold. Here we invoked the option to highlight certain chromosomes/scaffolds (specified in the configuration file; see Main text). Plot type 5 (**F**; see also Supplementary Figure 3 and 4) shows heterozygosity and genome coverage profiles for each studied individual and can be used to confirm that samples have been correctly sexed.

The first steps, trimming and subsampling of WGS reads (Figure 2a: Step 1), are optional. If the input reads are already trimmed for adaptors and bad quality sequences, this step can be skipped. The subsampling step may be used if the WGS files are unnecessarily large, in which case subsampling will reduce the pipeline run time. The reads are then aligned to the study-species reference genome (Step 2) and deduplicated (Step 3). From these deduplicated output BAM files (referred to as “unfiltered”), reads with different (and modifiable) number of mismatches to the reference genome are removed, and the remaining reads are written to new BAM files (Step 4). The default mismatch settings, which were used for all analyses in this paper, are: i) no filtering (“unfiltered”; see above), ii) intermediate filtering (≤2 mismatches allowed), and iii) strict filtering (0 mismatches allowed). Genome coverage is then calculated for each sample from these BAM files across 5 kb windows (Step 5). Inspecting the results from different mismatch settings is useful since the optimal mismatch threshold may differ between species and sex chromosome systems. Comparisons of different mismatch settings may also reveal important information about the level of sex chromosome differentiation, and the level of Y/W chromosome degeneration. The genome coverage values are normalized between samples (Step 6). Variants are called from the “unfiltered” BAM file (Step 7), followed by quality filtering (Step 8) and heterozygosity calculations across 5 kb windows for each sample (Step 9). Sex-specific genome coverage and heterozygosity mean values (if more than one sample per sex is used) is then calculated for each 5 kb window (Step 10). If the pipeline is run with the findZX snakefile, mean values will then be calculated over larger genome windows (Step 11), and output plots and tables will be generated (Step 12).

If the pipeline is run with findZX-synteny, a synteny analysis is performed between the two reference genomes (Step 13) and the genome coordinates from the study-species reference genome are lifted-over to the reference genome of a second species (Step 14). This option is recommended if the study-species reference genome is below chromosome-level, and/or to establish homology between sex chromosome systems (discussed below). The sex-specific genome coverage and heterozygosity values (Step 10) will then be translated to these liftover coordinates in the second species (Step 15). Lastly, the mean of these translated values will be calculated over larger genome windows (Step 16), and output plots and tables will be generated (Step 17).

Five types of output plots are generated by the pipeline (Figure 2b-f; see Supplementary Figure 1 for larger versions), all of which are discussed in detail below. If a list of chromosomes/scaffolds is provided as input to the pipeline, the plots will only show these ones. Similarly, certain chromosomes/scaffolds can be highlighted in plot type 4 if specified. All plotting steps are quick to re-run (e.g., with different sets of chromosomes/scaffolds and highlighting options) and do not require reanalysis of previous steps.

Four of the five output plot types (1-4; Figure 1b-e) are based on data from Step 11 (findZX) or 16 (findZX-synteny). In the last plot type (5; Figure 2f), genome coverage and heterozygosity values are plotted for each individual and mismatch setting separately (based on the data from Step 10 for findZX, Step 15 for findZX-synteny). Plot type 5 may be informative in confirming the sexing of included samples, and to inspect the genome coverage depth for each sample. Note, however, that for low differentiation sex chromosome systems, or systems where the sex chromosomes are extremely small, there may not be clear differences between sexes in these plots.

If the pipeline is run with one sample per sex together with a *de novo*-assembled study-species reference genome constructed from the homogametic sample, and if the species is highly heterozygous, the sex-specific signatures may be suboptimal and the results not clear. To circumvent this problem, the pipeline contains an option to construct a “consensus genome”, by incorporating genetic variants found in the two samples into the original study-species reference genome (Figure 2a; using the VCF file from Step 8; see also Methods). Running the pipeline again using this consensus reference genome instead of the original one has proven effective in a previous study, as it results in an autosome-wide equal mapping success between samples (for details see Sigeman et al. 2020). All output plots can be summarized in an interactive HTML report, with descriptions of each plot type.

### Datasets for pipeline evaluation

We evaluated the pipeline by analysing data from eight species with previously identified sex chromosomes (Supplementary Methods). These species were selected to represent a broad range of sex chromosome systems in terms of differentiation, size and age. Two species have highly differentiated sex chromosomes: platypus (*Ornithorhynchus anatinus*, multiple sex chromosome system; X_1-5_Y_1-5_; Grützner et al. 2004) and green anole lizard (*Anolis carolinensis*; XY; Alföldi et al. 2011). Three species have “neo-sex chromosomes” formed through fusions between the ancestral sex chromosomes and (parts of) autosomes. In these species, the added (formerly) autosomal regions are characterized by less sex chromosome differentiation than the old, ancestral part. The species with neo-sex chromosomes are: a fruit fly (*Drosophila miranda*; neo-XY; Zhou & Bachtrog 2012), Eurasian skylark (*Alauda arvensis*; neo-ZW; Sigeman et al. 2019; Dierickx et al. 2020), and mantled howler monkey (*Alouatta palliata*; neo-XY; Ma et al. 1975; Solari & Rahn 2005). Two species with low differentiation sex chromosome systems were also included: turquoise killifish (*Nothobranchius furzeri*; XY; Reichwald et al. 2015) and guppy (*Poecilia reticulata*; XY; Künstner et al. 2015; Wright et al. 2017). We also included the central bearded dragon (*Pogona vitticeps*; ZW), which has a micro-ZW sex chromosome system in which inferring homology to other species has proven to be challenging (but see Deakin et al. 2016). Lastly, we demonstrate the usefulness of the pipeline for identifying other divergent haplotype blocks using genomic data from different morphs of the ruff (*Calidris pugnax*), a bird species where male phenotypes (plumage colouration and behaviour) are determined by an inversion polymorphism (Lamichhaney et al. 2016; Supplementary Methods).

Accession numbers for all WGS data are provided in Supplementary Table 1, and accession numbers for all reference genomes used in Supplementary Table 2, and configuration files used to run each analysis is listed in Supplementary Table 3. The number of samples used for each species ranged between 1 and 3 individuals per sex, except for guppy where we ran the analyses with both a large (n = 23) and small (n = 2) number of individuals. For species with available chromosome-level assemblies (green anole lizard, fruit fly, turquoise killifish, platypus, and guppy), we ran the pipeline without a synteny-species (findZX). The remaining species were run with (and sometimes also without) a synteny-species (findZX-synteny). For several of the latter species, we used multiple different synteny-species. Supplementary Table 3 contains information on the number of samples used for each analysis, if the pipeline was run with (findZX-synteny) or without (findZX) a synteny-species reference genome, and what chromosomes/scaffolds were expected to be sex-linked/polymorphic (details regarding this are also described in the Supplementary Methods). Average genome coverage values for all BAM files (Figure 2a; Steps 3 and 4) are reported in Supplementary Table 4.

### Example output: mantled howler monkey

Here we show an example output from the analysis of one of the studied species, the mantled howler monkey. Previous studies have shown that this species has a neo-sex chromosome system, formed through one or two Y–autosome fusions (Ma et al. 1975; Solari & Rahn 2005). In a karyotype study including other *Alouatta* species, chromosome regions homologous to human chromosomes 3 and 15 were found to have fused to the ancestral Y chromosome (Consigliere et al. 1996). This species was chosen as an example output because its neo-sex chromosome system allows us to show both what highly differentiated regions (ancestral X; cf. Figure 1d) and more recently added regions of lower differentiation (fused formerly autosomal regions; cf. Figure 1b) look like in the pipeline output. We ran the pipeline using WGS data from 2 homogametic (XX) females and 2 heterogametic (XY) males, together with a newly published reference genome of this species (Supplementary Table 2), and both with and without the use of the human genome as a “synteny-species”. Plot types 1 (Figure 3), 3 (Figure 4a), and 4 (Figure 5c) are shown here in the Main text. Plot types 2 (Supplementary Figure 2) and 5 (Supplementary Figures 3 and 4) are in the Supplementary Information.

**Figure 3.**
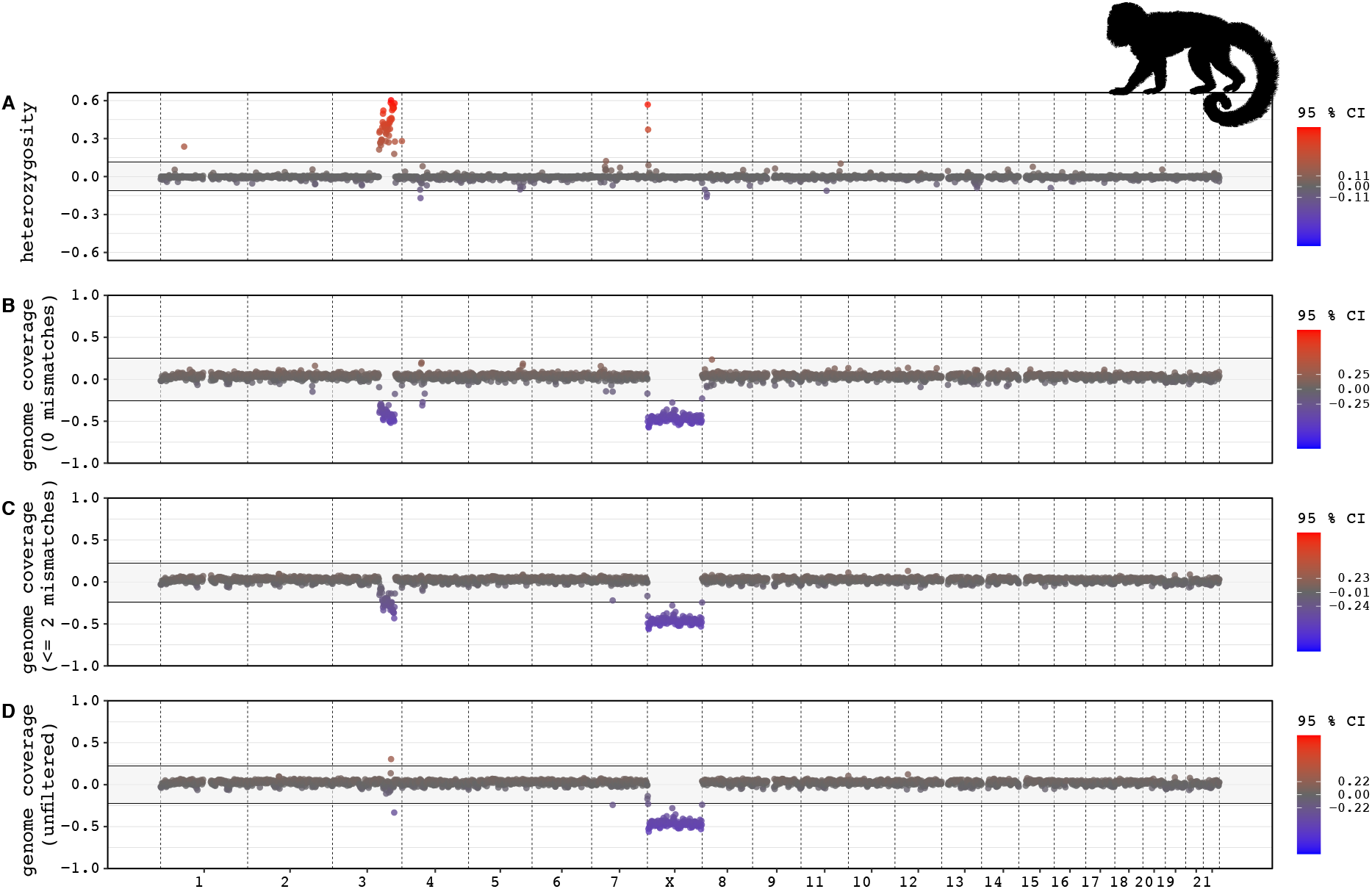
Sex differences in genome coverage and heterozygosity values (1 Mb windows) for the mantled howler monkeys, plotted along chromosome positions in the human genome. The four rows show: (**A**) heterozygosity, and genome coverage with (**B**) strict filtering (0 mismatches allowed), (**C**) intermediate filtering (≤2 mismatches) and (**D**) no filtering of mapped reads (“unfiltered”). The grey background marks the 95% confidence intervals (CI), and data points outside these values are red (if they are higher) or blue (if they are lower). All outlier values are reported as separate data tables in the pipeline output. The data reveals that chromosome X and a part of chromosome 3 are sex-linked in this species. The same data is also plotted for each sex separately (see Supplementary Figure 2).

**Figure 4.**
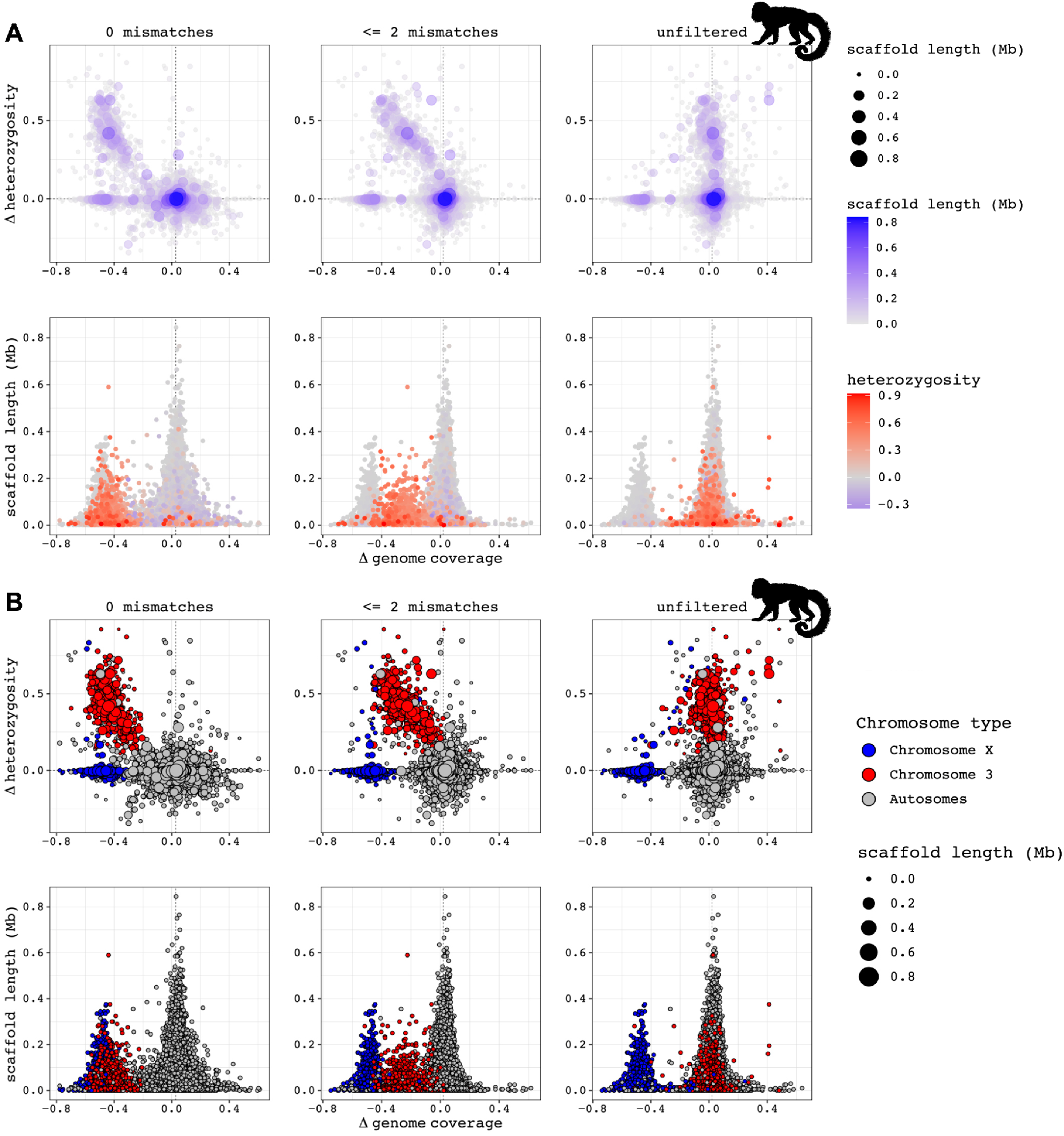
Per scaffold difference in heterozygosity and genome coverage between male and female mantled howler monkeys, with scaffold lengths indicated by symbol colour and size (top row in A and B) or plotted on the y-axis (bottom row in A and B). Genome coverage was calculated for three different mismatch filtering stringencies of mapped reads (left, mid, right panels: strict (0 mismatches), intermediate (≤2 mismatches) and no filtering (“unfiltered”). The left-side panels show three separate clusters of scaffolds, corresponding to the expected patterns of autosomes, and sex chromosome regions of low and high differentiation, respectively. (**A**) This plot was generated through the pipeline directly, with the findZX option (i.e., not using a synteny-species reference genome). (**B**) The underlying data in this plot is identical to Figure 4a but coloured differently to facilitate interpretation of Figure 4a (see Main text). In this plot, the data points are coloured blue if the scaffolds aligned to the human X chromosome and red if the scaffolds aligned to the sex-linked region of chromosome 3 (134-178 Mb; Figure 3). All other scaffolds are coloured grey.

**Figure 5.**
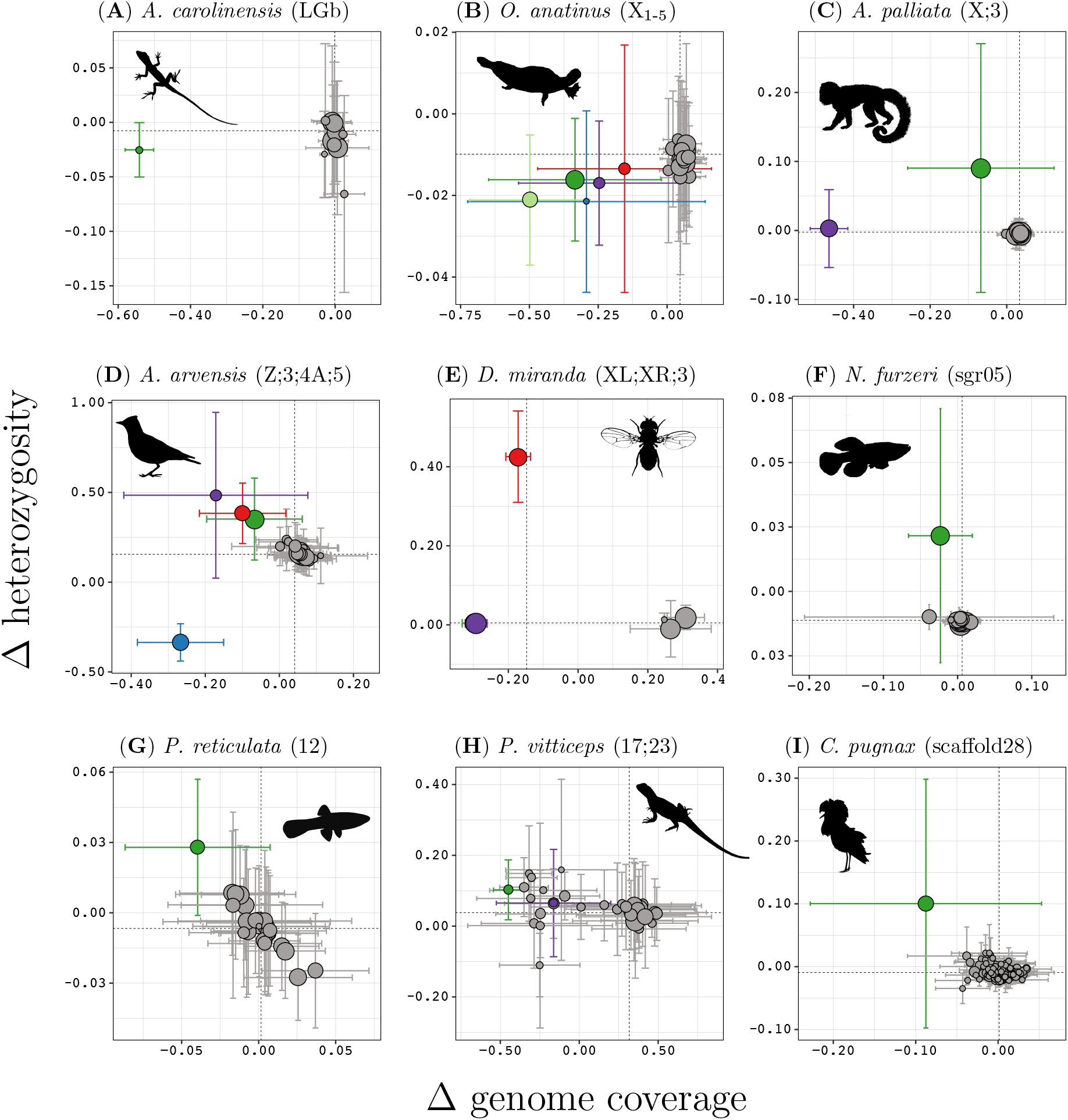
Genome coverage and heterozygosity values (mean ± standard deviation across 1 Mb windows) per chromosome or scaffold for all studied species. Dashed lines mark the genome-wide median across all 1 Mb windows. The chromosome/scaffold names in parentheses next to species names are the expected sex chromosomes/inversion polymorphism scaffolds as described in the Main text and Supplementary Information. These chromosomes/scaffolds are coloured differently from other chromosomes/scaffolds (which are grey). The sizes of the data points reflect the chromosome/scaffold length. Each of these plots constitutes one of 6 panels in Supplementary Figures 5-13 (shown here are the panels based on 0 mismatches, except for Eurasian skylark (*A. arvensis*) for which we show the panel based on ≤2 mismatches), which also have a colour legend for each species. These panels are based on pipeline runs using all samples listed for each species in Supplementary Table 1. All species except for mantled howler monkey (*A. palliata*), Eurasian skylark and central bearded dragon (*P. vitticeps*) were run using findZX. The mantled howler monkey was run, as described previously, with human (*H. sapiens*) as a synteny-species. Eurasian skylark was run with zebra finch (*T. guttata*) as a synteny-species, and for central bearded dragon we used the chicken (*G. gallus*) as a synteny-species. Silhouettes of animals are from phylopic (credits in Supplementary Information).

The reference genome of the mantled howler monkey consists of 43,519 scaffolds >10 kb and has a scaffold N50 value of 72 kb, meaning that scaffolds covering half the genome (1.5 Gb of the total size ~3 Gb) are shorter than 72 kb. Running the pipeline using a reference genome with this level of fragmentation is fine in principle (shown in Figure 4), but the use of a synteny-species (i.e., with findZX-synteny) with higher contiguity can add valuable information. The genome-wide plot types 1 (Figure 3; sex differences) and 2 (Supplementary Figure 2; values plotted for each sex separately) show that genomic regions in the mantled howler monkey with synteny to human chromosome X (i.e., the ancestral therian mammalian sex chromosome) and chromosome 3 have pronounced sex differences in heterozygosity and genome coverage compared to the rest of the chromosomes (Figure 3). The entire chromosome X (0-156 Mb) is characterized by markedly lower genome coverage in the heterogametic sex (male) compared to the homogametic sex (female), regardless of the number of allowed mismatches, concordant with the expected patterns from a heavily degenerated ancestral Y chromosome (homologous to chromosome X; Figure 3). Chromosome 3 shows characteristics of a younger and less degenerated sex-linked genomic region across parts of its length (134-178 Mb), as the genome coverage is reduced only when restricting the number of allowed mismatches and there is a clear excess of heterozygosity in the heterogametic sex compared to the homogametic sex (Figure 3).

The sex-specific signature across only parts of chromosome 3 may be the result of recombination suppression only extending to a part of the fused chromosome, or if only parts of chromosome 3 fused to the X chromosome. There was no sex-specific signature on chromosome 15, which has been shown to be fused to the Y chromosome in some *Alouatta* spp. (see above; Ma et al. 1975; Solari & Rahn 2005). This may be either because i) this chromosome is not fused to the rest of the sex chromosome in this species, or ii) that it is fused but recombination suppression has not extended to this part of the sex chromosomes.

There are several reasons for running the pipeline without the use of a synteny-species (i.e., with findZX): (1) when the study-species reference genome itself is of sufficiently high contiguity for the purposes of the study, (2) when the purpose is to identify sex-linked scaffolds in the study-species reference genome, or (3) when no suitable synteny-species reference genome is available. For highly contiguous study-species reference genomes, output plot types 1 (Figure 3) and 2 (Supplementary Figure 2) may reveal large coherent sex-linked regions. However, in cases where the study-species reference genome is highly fragmented (as in the mantled howler monkey), plot type 3 (Figure 4) is more likely to be of use. This plot type displays differences in sex-specific heterozygosity and genome coverage values per scaffold (as opposed to across large genome windows), while also separating the scaffolds by length (Figure 4a). The latter is valuable as short scaffolds may be responsible for much noise in the data.

To facilitate interpretation of plot type 3 (Figure 4a), we produced a second plot (Figure 4b) using the same data but where the scaffolds in the mantled howler monkey genome aligning to sex-linked regions identified when using human as synteny-species (Figure 3) were coloured differently from the rest. Specifically, we coloured all scaffolds from the mantled howler monkey genome aligning with more than 90% of their length to the X chromosome (0-156 Mb) in blue, and those aligning to the sex-linked part of chromosome 3 (134-178 Mb) in red. All other scaffolds were coloured grey. Three clusters are clearly defined in the leftmost panels of Figure 4b corresponding to different chromosomal categories (cf. Figure 1): one (grey) cluster with no or little differentiation between sexes (autosomes), one (blue) with a slight deficiency of heterozygous sites and low genome coverage in the heterogametic sex (highly differentiated sex chromosomes), and one (red) with an excess of heterozygosity in the heterogametic sex and lower genome coverage in the heterogametic sex (low differentiation sex chromosomes). The effect of restricting the allowed number of mismatches between mapped reads and the reference genome for low differentiation sex-linked regions can be clearly seen in Figure 4, as a cluster of scaffolds with an excess of heterozygosity and lower genome coverage in the heterogametic sex (red cluster in Figure 4b) “moves” towards the autosomal cluster when allowing reads with more mismatches to be included (from left-side panel to right-side panel).

### Validation through analysis of nine datasets

To analyse the performance of this pipeline for sex chromosome systems of varying sizes and degrees of differentiation, we applied it to downloaded datasets for all the previously listed species (n = 9; Supplementary Table 1). Plots supporting results that are specifically mentioned in the Main text are provided as Supplementary Figures 5-22. In this section, we show excerpts of output plot type 4 (Figure 5; see the full six-panel plots in Supplementary Figures 5-13). The underlying data for plot type 4 is the same as for plot type 1 (Figure 3) and 2 (Supplementary Figure 2) but is in the format of scatter plots (with heterozygosity on the y-axis and genome coverage on the x-axis) instead of genome-wide plots (with chromosome position on the x-axis). The mean (± standard deviation) values for all genome windows per chromosome (here 1 Mb windows) are shown in Figure 5. We selected the previously identified sex chromosome(s) to be distinguished from other chromosomes or scaffolds with the “highlighting” option (see above). In the case of the ruff, the scaffold previously found to contain the 4.5 Mb large inversion polymorphism was highlighted (Lamichhaney et al. 2016). Details on what chromosomes or scaffolds are expected to be sex-linked (or associated with male morphs) are in the Supplementary Information (and summarized in Supplementary Table 3).

In eight of the nine species, the expected chromosome(s)/scaffold(s) were clearly distinguished from the rest of the chromosomes/scaffolds (Figure 5a-g,i). Among these were both the species with high sex chromosome differentiation and Y degeneration: the green anole lizard (chromosome LGb; Figure 5a; Supplementary Figure 5) and the platypus (chromosomes X_1_-X_5_; Figure 5b; Supplementary Figure 6). The sex chromosomes of these species were identified through lower genome coverage in the heterogametic (XY) sex, which also had lower or similar levels of heterozygosity (Figure 5a,b). The three neo-sex chromosome systems were also clearly identified: the mantled howler monkey (chromosomes X, 3; Figure 5c; Supplementary Figure 7), the Eurasian skylark (chromosomes Z, 3, 4A, 5; Figure 5d; Supplementary Figure 8) and the fruit fly (chromosomes XL, XR and 3; Figure 5e; Supplementary Figure 9). The sex chromosomes in these species consist of an ancestral region of high differentiation and Y/W degeneration as well as regions of more recent sex-linkage, which are characterized by less differentiation and Y/W degeneration. The ancestral sex chromosome regions in these species were identified through lower genome coverage and equal to lower heterozygosity in the heterogametic sex (mantled howler monkey: chromosome X, Figure 5c; Eurasian skylark: chromosome Z, Figure 5d; fruit fly: chromosome arms XL and XR, Figure 5e). The added regions were characterized by higher heterozygosity and similar-to-lower genome coverage in the heterogametic sex (mantled howler monkey: chromosome 3, Figure 5c; Eurasian skylark: chromosomes 3, 4A and 5, Figure 5d; fruit fly: chromosome 3, Figure 5e). Note that the genome coverage values for the autosomes in the fruit fly are >0.2 and that the values from the sex chromosome ranges between −0.15 to −0.25, while the autosomes in the other species have values around 0. This difference can be explained by the sex chromosomes constituting a very large part of the genome in the fruit fly, which affects the normalization between sexes.

Three species, turquoise killifish, guppy, and the central bearded dragon (Figure 5f-h; Supplementary Figure 10-12), have sex chromosome systems that have been challenging to identify, due to either low differentiation (turquoise killifish and guppy) or due to difficulties in establishing homology to other species (central bearded dragon). Of these, we successfully identified the turquoise killifish (sgr05; Figure 5f) and the guppy (chromosome 12; Figure 5g) sex chromosomes. The expected sex chromosome regions were not clear outliers in the central bearded dragon (chromosomes 17 and 23; Figure 5h). However, chromosome 17 (Figure 5h, in green) did have the lowest male-to-female genome coverage of all chromosomes. Lastly, the scaffold containing the inversion polymorphism controlling male morphs in the ruff (scaffold28/NW_015090842.1) was identified through having lower genome coverage and higher heterozygosity in the “faeder” phenotype individual (which is heterozygotic for the inversion) than in the “resident” phenotype individuals (which is homozygotic for the ancestral, non-inverted haplotype; Figure 5i; Supplementary Figure 13).

### Robustness to evolutionary distances, sample sizes and genome coverage

To test for the effect of lowering the sample size for species whose sex chromosomes had been successfully identified based on more than one sample per sex, i.e. mantled howler monkey, turquoise killifish and guppy (Figure 5c,f,g), we reran the pipeline using only 1 individual per sex. In all three cases, the same regions were retrieved (Supplementary Figures 14-16). To test for sensitivity to sequencing depth, we subsampled the WGS data from 1 female and 1 male mantled howler monkey to 50% of the number of base pairs in the smallest of the mantled howler monkey samples. This did not affect the results either, as the same sex-linked regions were retrieved, even with as low as 1.43 X coverage with 0 mismatches allowed (Supplementary Figures 17; average genome coverage values for all discussed analyses are given in Supplementary Table 4).

The results from the mantled howler monkey with human as synteny-species (Figure 3) demonstrate that the pipeline is robust to large evolutionary distances between the studied species and the synteny-species, as howler monkeys (that belong to the New World monkeys) and humans separated over 35 million years ago (Vanderpool et al. 2020). We also ran the pipeline with an even more distant relative to the mantled howler monkey as a synteny-species; the meerkat (*Suricata suricatta*; Supplementary Table 2; Dudchenko et al. 2017; Dudchenko et al. 2018). Even though these species are separated by over 80 million years of independent evolution (dos Reis et al. 2012), we still managed to identify the sex-linked regions (homologous to meerkat chromosomes X and 5; Supplementary Figure 18). Additionally, we ran the Eurasian skylark samples with the chicken (*Gallus gallus*) as a synteny-species reference genome and obtained the expected sex-linked regions (Supplementary Figure 19 and 20). Lastly, we ran the ruff samples using the chicken as a synteny-species and found chromosome 11 to be an outlier, as predicted (Supplementary Information; Supplementary Figure 21). Both the Eurasian skylark and the ruff separated from the chicken over 70 million years ago (Prum et al. 2015).

## Discussion

We present a novel Snakemake pipeline and demonstrate its effectiveness in uncovering sex-linked genomic regions using WGS data from a wide range of species and sex chromosome systems. The pipeline is easy to use, and the output plots can be easily customized in terms of what data is shown, thereby producing near-publication ready figures. The findZX-synteny option allows the user to quickly infer homology between sex chromosomes in different species, and we show that this method is effective even when the study-species and synteny-species are phylogenetically distant (Supplementary Figures 18-21). All heterozygosity and genome coverage values used to produce the figures, as well as outlier values outside the 95% CI genome-wide averages based on the specified window sizes (from plot type 1), are output as separate tables which facilitates further data exploration. A README file in the table directory describes each column, and names of output tables used for plotting are specified in each output plot.

Sex chromosomes can be identified using different kinds of genomic data (Muyle et al. 2016; Rangavittal et al. 2019; Feron et al. 2020; Palmer et al. 2019). Here we utilize WGS data, which is more comprehensive compared to reduced representation sequencing techniques (such as RADseq) or RNA-seq, thereby increasing the likelihood of sex chromosome identification. It also provides opportunities for follow-up studies of e.g., gene content and repeat landscape of sex-linked regions. WGS sequencing is more expensive per sample than for example RADseq. However, the results from the multi-species analyses (Figure 5) show that one sample per sex is sufficient to reveal sex-linked regions in most systems (Figure 5; Supplementary Table 3). We also show that the method is robust to low sequencing coverage (Supplementary Table 4). Note, however, that too low coverage will lead to an underestimation of heterozygous sites. This pipeline cannot be directly compared to other published methods and software for detecting sex chromosomes, as it uses a different data type. The availability of complementary methods for identifying sex chromosomes is highly useful, as it allows the scientific community to take full advantage of the variety of sequence data that are being produced.

Our pipeline successfully detected the previously identified sex-linked regions in seven of the eight studied species (Figure 5a-f). One of these species was the guppy, in which the sex chromosomes have been notoriously difficult to characterise, even when analysing many samples per sex (Wright et al. 2017). We were also able to identify the inversion polymorphism present in the ruff (Figure 5i). However, we found no clear sex-linkage signature for the expected chromosomes in the central bearded dragon (Figure 5h). Note also that when the fully sex-linked region is very small (as e.g., in the extreme example of the fugu; Kamiya et al. 2012), our approach is likely too crude as it is based on genome window scans, and per-chromosome or -scaffold calculations. Our code is, however, open and modifiable for any special needs.

We urge users to inspect all the different output plots to avoid misinterpretations. For example, our method assumes that the type of heterogamety (XY or ZW) is known a priori. Should a heterogametic study-system reference genome be used instead of a homogametic one, one sex will show an excess of genomic windows without any (or with very low) coverage in the other sex (representing Y/W scaffolds). Plot type 5, which shows per-individual genome coverage and heterozygosity profiles (Supplementary Figures 3 and 4) may reveal if any of the included samples were not sexed correctly, and the pipeline can be rerun using correct heterogamety settings for these samples. It is also important to look not only at the female-to-male difference values (Figure 3) when interpreting the results, but also the values for the different sexes separately (Supplementary Figure 2). A low heterogametic-to-homogametic genome coverage signature may be caused by either lower coverage in the samples of the heterogametic sex, or by heightened coverage values in the samples of the homogametic sex. An example of the latter seems to be sgr13 in the turquoise killifish, in which the homogametic sample appears to have higher than genome-average coverage while the heterogametic sample has genome coverage similar to the genome-average (Supplementary Figure 22). In this case, there is no difference in heterozygosity compared to the genome average in either sex, which would be expected if this region was truly sex-linked. This may result from the presence of a duplicated region in the homogametic individual (i.e., structural variation occurring in the population) compared to the reference genome, although more samples would be needed to confirm this hypothesis. Independent validation of sex-linked regions found using this pipeline is recommended, for example through PCR and Sanger sequencing of targeted loci in additional samples, especially when analysing only one sample of each sex (cf. Sigeman et al. 2020).

Uncovering novel changes in sex chromosome systems between species is important to advance our understanding of sex chromosome evolution. Furthermore, failing to consider unexpected sex chromosome variation may lead to misinterpretations of genomic patterns in studies not related to sex chromosome research. As WGS data becomes increasingly available (and published on open databases such as NCBI’s short read archive), we hope that our pipeline will be of use to many researchers.

## Methods

Information on how to install findZX is in the Supplementary Information and on the GitHub page (https://github.com/hsigeman/findZX). All needed software and dependencies, which are fully compatible with both Linux (64-bit) and macOS systems, can be easily installed using the Conda package manager (Anaconda Software Distribution, 2016). The minimal data requirements for running findZX is paired-end WGS data from one individual of each sex, as well as a reference genome of the homogametic sex from the same species (Figure 2a). If no reference genome is available for the study-species, it can be constructed based on the paired-end data from the homogametic sample (see instructions on the GitHub page). To run findZX-synteny, a reference genome of the species that should be used for the genome coordinate liftover analysis is also required (Figure 2a). The analysis will be run according to settings provided in a configuration file, which also contains paths to input data (see the GitHub page and Supplementary Information for usage examples). The following sections describe the different computational steps performed by findZX.

### Trimming and subsampling (optional)

The first step performed by findZX, if opted for in the configuration file, is trimming and quality filtering of the fastq reads using Trimmomatic v0.39 (Bolger et al. 2014). An HTML quality control report will be created for both the untrimmed and trimmed reads, using fastqc v0.11.9 (Andrews 2010) and multiqc v1.10.1 (Ewels et al. 2016). Very large fastq files can take a long time to process (especially the alignment of reads to the reference genome). FindZX therefore includes an option to subsample reads to a specific number of basepairs using reformat.sh (BBTools suite v38.92, Bushnell). The pipeline is written in a modular way so that it can be stopped after the trimming and HTML report are completed, to allow for visual inspection of the results before running the rest of the steps. FindZX can then be restarted if the trimming was successful.

### Alignment and variant calling

The reads from each sample are then aligned to the reference genome of the study species using bwa mem v0.7.17 (Li & Durbin 2009a), sorted using samtools sort v1.7 (Li et al. 2009b) and deduplicated with picardtools v2.18.0 (http://broadinstitute.github.io/picard). Variants are called for all samples using platypus v0.8.1.1 (Rimmer et al. 2014) and compressed using bgzip and tabix v1.9 (Li 2011).

### Heterozygosity and genome coverage calculations

Variants are filtered with vcftools v0.1.15 (Danecek et al. 2011) using options –gzip –min-alleles 2 – max-alleles 2 –minQ 20 –MinDP 3 and –remove-filtered-geno-all. The number of heterozygous sites is summed across 5 kb genome windows for each individual. For each genome window, the mean proportion of heterozygous sites (number of heterozygous sites per 100 base pairs) is calculated for each sex.

Genome coverage is calculated using bedtools multicov v2.27.1 (Quinlan & Hall, 2010) (only considering properly aligned reads with quality score >20) across the same 5 kb genome windows. This is done on the original deduplicated BAM files (see above) which will be referred to as “unfiltered”, and on two other BAM files that are filtered to only contain reads with different thresholds of mismatches to the reference genome (thresholds specified in the configuration file). The thresholds used for all analyses in this paper are i) “unfiltered”, ii) intermediate filtering (≤2 mismatches allowed), and iii) strict filtering (0 mismatches allowed). The genome coverage values for each sample in each of these three BAM files are first filtered for outliers (values exceeding the genome-wide mean ± three times the standard deviation are masked), and then normalized between samples. Alignment statistics for each BAM file are generated using samtools stats v1.7.

### Chromosome anchoring to a synteny-species (only when using findZX-synteny)

If findZX-synteny is used, a lastdb (Kiełbasa et al. 2011) database will be constructed from the synteny-species reference genome. Whole-genome alignment between the study-species and synteny-species reference genomes (using the lastdb database) will then be performed with lastal, and converted to psl format (using maf-convert); both programs from the software last v1238 (Kiełbasa et al. 2011). A custom script then locates the longest match for each 5 kb window in the study-species reference genome to the synteny-species reference genome, with 500 matching base pairs as a minimum. Matches where more than two windows from the study-species matched to the same chromosome position in the synteny-species reference genome are filtered out.

### Statistics, plotting and HTML report

Mean heterozygosity (from the “unfiltered” BAM file) and genome coverage values (from BAM files with all three mismatch filtering settings; see above) are calculated per sex and chromosome/scaffold, and per sex across different genome window sizes. The window sizes are specified in the configuration file (for the analyses in this paper we chose 50 kb, 100 kb and 1 Mb for all species). From these values, differences between sexes are calculated and written as output tables. Genome-wide mean ± 95% confidence intervals (CI) are calculated for the heterozygosity and genome coverage data, and another set of tables is written with windows having values on either side of these thresholds (mean ± 95% CI).

The sex differences in heterozygosity and genome coverage are plotted in different formats (see Results section and Supplementary Information for examples) per chromosome/scaffold, and per window size. If a list of chromosomes/scaffolds is provided in the configuration file, these are the only ones that will be plotted. If no such file is provided all chromosomes/scaffolds will be plotted, except for the genome-wide plots (plot type 1 and 2) which require a pre-set maximum number of scaffolds (default: n = 50). FindZX also provides output plots that can be used for verifying the sex of the studied samples by plotting heterozygosity and genome coverage profiles for each sample separately (plot type 5). Note, however, that these results are clearer for species with large and heteromorphic sex chromosomes.

All output plots are in multi-page PDF format, where the last page contain information on what output table was used to create each plot. All output plots can be rendered into an interactive HTML report file, which will also contain run time information and the multiqc report files (see Supplementary Information/GitHub page for details).

### Consensus genome assembly

Results from a previous study showed that sex chromosome regions of low differentiation (Figure 1a) can be hard to identify in highly heterozygous species, when only one sample of each sex is used and when the reference genome is constructed from one of these samples (Sigeman et al. 2019). This is because identification of such regions often requires strict filtering of mismatches to the reference genome (Figure 4), and in highly heterozygous species this may lead to a drastic reduction of aligned reads both from the autosomes and the sex chromosomes in the heterogametic sample which will obscure the signal from the sex chromosomes (see Sigeman et al. 2019 for more details). To circumvent this problem, findZX has an option to create a consensus reference genome using bcftools consensus, in which all biallelic variants with non-reference allele counts ≥2 will be incorporated. The consensus reference genome is created from a filtered version of the output VCF file (Figure 2a; Step 8). This new consensus genome assembly can then be used as input to the pipeline instead of the original reference genome. See Supplementary Information and/or the GitHub page for instructions on how to run this.

### Per-sample average genome coverage calculations

We performed one analysis in this paper that was not part of the pipeline: calculations of an average genome coverage value per sample (Supplementary Table 4). These calculations were done using a bash script (available on GitHub at workflow/scripts/calc_cov.sh) which takes as input the output file from samtools stats (see above) and the indexed study-species reference genome (produced with samtools faidx). The script divides the total number of aligned base pairs in each sample by the total length of the respective study-species reference genome.

## Availability of data and materials

The findZX pipeline is available on GitHub: https://github.com/hsigeman/findZX, along with instructions on how to install and configure findZX for analyses on new datasets. A short tutorial is also provided in the Supplementary Information. All sequencing data and reference genomes used in this study are publicly available and listed in Supplementary Tables 1 and 2. Paths to configuration files for all findZX analyses are also given in Supplementary Table 3.

## Acknowledgements

Bioinformatics analyses were performed on computational infrastructure provided by the Swedish National Infrastructure for Computing (SNIC) at Uppsala Multidisciplinary Center for Advanced Computational Science (UPPMAX). The research was funded by grants from Vetenskapsrådet (Consolidator Grant No. 2016-00689 to B.H). We thank Verena E Kutschera for suggestions on how to improve the pipeline, and for comments on the manuscript. We also thank the SexGen research group at Lund University for their help with software testing.

## Notes

https://github.com/hsigeman/findZX

### Competing Interest Statement

The authors have declared no competing interest.

