## Supplementary Information for "FindZX: an automated pipeline for detecting and visualising sex chromosomes using whole-genome sequencing data"

### 1 Supplementary Methods

#### 1.1 How to install and run findZX

##### 1.1.1 Download and installation

FindZX works on Linux and macOS systems, and contains a configuration file which can be used to run the pipeline on a SLURM system. There is only one prerequisite (except for findZX itself): that conda (`https://docs.conda.io/en/latest/`) is installed on the system. Conda will download and install all other dependencies into separate conda environments the first time findZX/findZX-synteny is run.

Obtain a copy of findZX from the GitHub page:

```
git clone https://github.com/hsigeman/findZX.git
cd findZX # Go to directory
```

Create a minimal conda environment and install software automatically through findZX:

```
conda create -n findZX -c conda-forge -c bioconda \
python=3.9.4 snakemake-wrapper-utils=0.2.0 snakemake=6.4.0 \
mamba=0.15.3
```

Then activate the environment:

```
conda activate findZX
```

To verify the installation, run findZX on a test dataset:

```
snakemake -s workflow/findZX --configfile .test/config.yml \
--cores 1 -R all --use-conda -k
```

For more information, please visit the GitHub page (`https://github.com/hsigeman/findZX.git`).

##### 1.1.2 Basic usage

FindZX can be run using the following command:

```
snakemake -s workflow/findZX \
--configfile <Config file> \
--cores <Select number of cores> -R all \
--use-conda -k
```

FindZX-synteny can be run using the following command:

```
snakemake -s workflow/findZX-synteny \
--configfile <Config file> \
--cores <Select number of cores> -R all \
--use-conda -k
```

Note that computational steps shared by findZX and findZX-synteny (Figure 2a in the Main text; Steps 1-10) do not need to be rerun when changing between using findZX and findZX-synteny. If an analysis is first performed using findZX, and later using findZX-synteny (adding a synteny-species), only Steps 13-17 are run. Similarly, if an analysis is first performed using findZX-synteny and then findZX, only Steps 11-12 are run. This is because snakemake automatically keeps track of which files need to be created, and which ones are already present.

To render an interactivate HTML report with all output plots and run times, run the following command once findZX/findZX-synteny is finished:

```
snakemake -s workflow/findZX{-synteny} \
--configfile <Config file> \
--report <Report_name.html>
```

The pipeline can also be run on a SLURM server cluster, by specifying a SLURM configuration file (cluster.yml):

```
snakemake -s workflow/findZX \
--configfile <Config file> \
--cores <Select number of cores> -R all \
--cluster-config cluster.yml \
--cluster " sbatch -A {cluster.account} -t {cluster.time} \
-n {cluster.n} " \
--use-conda -k
```

For more information on how to use findZX/findZX-synteny, see instructions on the GitHub page (`https://github.com/hsigeman/findZX`)

##### 1.1.3 Constructing a consensus genome

Use this command to run the ”consensus genome” option (see Main text):

```
snakemake -s workflow/findZX \
--configfile <Config file> \
--cores <Select number of cores> -R modify_genome \
--use-conda -k
```

See below (*Alauda arvensis* analysis under subsection 1.2.1) for example usage.

##### 1.1.4 Reproducing the results in this paper

The commands used to run all analyses are described for each species under subsection 1.2, and summarized in Supplementary Table 3.

#### 1.2 Information on the sex chromosome systems of the studied species, and details on findZX analyses:

Here, we give brief information about the current knowledge of the sex chromosome system in each of the studied species (or inversion polymorphism in the case of the ruff), and how each species was analysed. In species where a synteny-species reference genome was used to anchor the scaffolds/chromosomes from the study-species reference genome, we list which chromosomes are expected to show signs of sex-linkage in each synteny-species. These expectations are based either on results from previous studies or based on known syntenies between species (as determined by https://www.genomicus.biologie.ens.fr/).

Config files listed here are on the findZX GitHub page.

##### 1.2.1 *Alauda arvensis* (Eurasian skylark)

**Sex chromosome system:**
Neo-sex chromosome system with shared synteny to zebra finch (*Taeniopygia guttata*) chromosomes Z, 3, 4A and 5 [1, 2]

**Details on findZX analyses:**
The Eurasian skylark WGS samples (n = 2; Supplementary Table 1) were analysed using a (highly fragmented) study-species reference genome constructed from the male WGS sample (Supplementary Table 1, Supplementary Table 3; see Sigeman et al. [1]).

We ran the findZX pipeline three times for this species. First, we used the "consensus reference genome" option, to ensure equal mapping success to the reference genome between sexes (with findZX):

```
snakemake -s workflow/findZX --configfile \
config/9_species_config/Alauda_arvensis_config.yml \
--cores 25 -R modify_genome --use-conda -k
```

Then, we ran the pipeline using this consensus reference genome as input to the pipeline, using the zebra finch (*Taeniopygia guttata*) as a synteny-species (with findZX-synteny):

```
snakemake -s workflow/findZX-synteny --configfile \
config/9_species_config/Alauda_arvensis_consensus_config_ZF.yml \
--cores 25 -R all --use-conda -k
```

And lastly using the chicken (*Gallus gallus*) as a synteny-species (with findZX-synteny):

```
snakemake -s workflow/findZX-synteny --configfile \
config/9_species_config/Alauda_arvensis_consensus_config_GG.yml \
--cores 25 -R all --use-conda -k
```

**Expected sex-linked regions:**
***Synteny-species 1 (zebra finch):*** chromosomes Z, 3, 4A and 5 [1, 2].
***Synteny-species 2: (chicken):*** chromosomes Z, 3, 4 and 5 (Based on synteny between zebra finch and chicken genomes: https://www.genomicus.biologie.ens.fr/.

##### 1.2.2 *Alouatta palliata* (mantled howler monkey)

**Sex chromosome system:**
Multiple sex chromosome system ($X_1X_2Y$) [3]. The $X_2$ chromosome in *A. caraya* (a relative of *A. palliata*) share synteny with human chromosomes 3 and 15 [4].

**Details on findZX analyses:**
The mantled howler monkey WGS samples (n = 4; Supplementary Table 1) were analysed using

a fragmented study-species reference genome (Supplementary Table 3). We first ran the pipeline using all samples (n = 4) and with the findZX option:

```
snakemake -s workflow/findZX --configfile \
config/9_species_config/Alouatta_palliata_config.yml \
--cores 25 -R all --use-conda -k
```

Then, we used the human (*Homo sapiens*) reference genome (Supplementary Table 3) as a synteny-species (findZX-synteny):

```
snakemake -s workflow/findZX-synteny --configfile \
config/9_species_config/Alouatta_palliata_config.yml \
--cores 25 -R all --use-conda -k
```

We then ran the pipeline (again with human as a synteny-species) using only one sample of each sex (n = 2):

```
snakemake -s workflow/findZX-synteny --configfile \
config/9_species_config/Alouatta_palliata_config_1M_1F.yml \
--cores 25 -R all --use-conda -k
```

Then with downsampling (to 50 % of the basepairs in the sample with the smallest number of reads) of the same two samples (n = 2):

```
snakemake -s workflow/findZX-synteny --configfile \
config/9_species_config/Alouatta_palliata_config_subsampling_1M_1F.yml \
--cores 25 -R all --use-conda -k
```

Then, lastly using the meerkat (*Suricata suricatta*) (Supplementary Table 3) as synteny-species reference genome (findZX-synteny):

```
snakemake -s workflow/findZX-synteny --configfile \
config/9_species_config/Alouatta_palliata_config_SS.yml \
--cores 25 -R all --use-conda -k
```

**Expected sex-linked regions:**
***Study-species:*** Unknown
***Synteny-species 1 (human):*** chromosomes X, 3, 15 [4].
***Synteny-species 2: (meerkat):*** chromosomes X, 5, 9 (Based on synteny between human and meerkat genomes: https://www.genomicus.biologie.ens.fr/.

##### 1.2.3 *Anolis carolinensis* (green anole)

**Sex chromosome system:**
Small XY-system, chromosome LGb [5].

**Details on findZX analyses:**
The green anole WGS samples (n = 2; Supplementary Table 1) were analysed using the chromosome-level reference genome of this species (Supplementary Table 3) as a study-species reference genome (with findZX):

```
snakemake -s workflow/findZX --configfile \
config/9_species_config/Anolis_carolinensis_config.yml \
--cores 25 -R all --use-conda -k
```

**Expected sex-linked regions:**
***Study-species:*** chromosome LGb [5].

###### 1.2.4 *Calidris pugnax* (ruff)

**Inversion polymorphism:**
Scaffold28 contains the 4.5 Mb large inversion polymorphism controlling male phenotypes in the ruff [6]. Scaffold28 shares synteny with chicken chromosome 11 [6].

**Details on findZX analyses:**
The ruff samples (n = 3; Supplementary Table 1) were analysed using the scaffold-level reference genome of the ruff (Supplementary Table 3) as a study-species reference genome. The pipeline was run twice; first without a synteny-species reference genome (findZX):

```
snakemake -s workflow/findZX --configfile \
config/9_species_config/Calidris_pugnax_config.yml \
--cores 25 -R all --use-conda -k
```

And then using the chicken as a synteny-species reference genome (findZX-synteny):

```
snakemake -s workflow/findZX-synteny --configfile \
config/9_species_config/Calidris_pugnax_config.yml \
--cores 25 -R all --use-conda -k
```

**Expected inversion polymorphic region:**
***Study-species:*** scaffold28/NW_015090842.1 [6].
***Synteny-species (chicken):*** chromosome 11 [6].

###### 1.2.5 *Drosophila miranda* (fruit fly)

**Sex chromosome system:**
Neo-XY system involving chromosomes XL, XR (ancestral sex chromosome arms) and Muller-C (fusion) [7].

**Details on findZX analyses:**
The fruit fly samples (n = 2; Supplementary Table 1) were analysed using the chromosome-level reference genome of this species (Supplementary Table 3) as a study-species reference genome (findZX):

```
snakemake -s workflow/findZX --configfile \
config/9_species_config/Drosophila_miranda_config.yml \
--cores 25 -R all --use-conda -k
```

**Expected sex-linked regions:**
***Study-species:*** Chromosomes XL (NC_030302.1), XR (NC_030303.1) and Muller-C (chromosome 3; NC_030305.1) [7].

###### 1.2.6 *Nothobranchius furzeri* (turquoise killifish)

**Sex chromosome system:**
Small relatively undifferentiated XY-system [8].

**Details on findZX analyses:**
The turquoise killifish WGS samples (n = 3; Supplementary Table 1) were analysed using a chromosome-level study-species reference genome (Supplementary Table 3). We first ran the pipeline (findZX) using all samples (n = 3):

```
snakemake -s workflow/findZX --configfile \
config/9_species_config/Nothobranchius_furzeri_config.yml \
--cores 25 -R all --use-conda -k
```

And then using only one sample of each sex (n = 2).

```
snakemake -s workflow/findZX --configfile \
config/9_species_config/Nothobranchius_furzeri_config_1M_1F.yml \
--cores 25 -R all --use-conda -k
```

**Expected sex-linked regions:**
***Study-species:*** Chromosome sgr05 [8].

##### 1.2.7 *Ornithorhynchus anatinus* (platypus)

**Sex chromosome system:**
XY-system consisting of five X chromosomes and five Y chromosomes: $X_{1\text{-}5}Y_{1\text{-}5}$ [9].

**Details on findZX analyses:**
The platypus WGS samples (n = 2; Supplementary Table 1) were analysed (findZX) using a chromosome-level study-species reference genome (Supplementary Table 3):

```
snakemake -s workflow/findZX --configfile \
config/9_species_config/Ornithorhynchus_anatinus_config.yml \
--cores 25 -R all --use-conda -k
```

**Expected sex-linked regions:**
***Study-species:*** Chromosome $X_1$-$X_5$ [9].

##### 1.2.8 *Poecilia reticulata* (guppy):

**Sex chromosome system:**
Extremely undifferentiated XY-system [10, 11], LG12/chr12.

**Details on findZX analyses:**
All guppy WGS samples (n = 23; Supplementary Table 1) were analysed (findZX) using a chromosome-level study-species reference genome (Supplementary Table 3):

```
snakemake -s workflow/findZX --configfile \
config/9_species_config/Poecilia_reticulata_config.yml \
--cores 25 -R all --use-conda -k
```

And then using two (n= 2) of the original WGS samples:

```
snakemake -s workflow/findZX --configfile \
config/9_species_config/Poecilia_reticulata_config_1M_1F.yml \
--cores 25 -R all --use-conda -k
```

**Expected sex-linked regions:**
***Study-species:*** Chromosome 12/LG12 [10, 11].

##### 1.2.9 *Pogona vitticeps* (central bearded dragon)

**Sex chromosome system:**
Micro-ZW system. Share partial synteny with chicken microchromosomes 17 and 23 [12].

**Details on findZX analyses:**
The central bearded dragon samples (n = 6; originating from 2 individuals; Supplementary Table 1) were analysed using the scaffold-level reference genome of the central bearded dragon (Supplementary Table 3) as a study-species reference genome. The pipeline was run using the chicken as a synteny-species reference genome (findZX-synteny):

```
snakemake -s workflow/findZX-synteny --configfile \
config/9_species_config/Pogona_vitticeps_config.yml \
--cores 25 -R all --use-conda -k
```

**Expected sex-linked regions:**
***Synteny-species (chicken):*** Chromosome 23 and 17 [12].

### 2 Supplementary Tables

Supplementary Table 1: WGS paired-end data used for analyses. SRA files from the same BioSample (only relevant for *Pogona vitticeps*) were treated as different samples.

| Species | Common name | Sex | Heterogametic/homogametic | SRA code | BioSample |
|---|---|---|---|---|---|
| *Alauda arvensis* | Eurasian skylark | female | Heterogametic | SRR10340221 | SAMN13107494 |
| *Alauda arvensis* | Eurasian skylark | male | Homogametic | SRR10340220 | SAMN13107495 |
| *Alouatta palliata* | mantled howler monkey | male | Heterogametic | SRR9655170 | SAMN10407285 |
| *Alouatta palliata* | mantled howler monkey | male | Heterogametic | SRR9655171 | SAMN10407294 |
| *Alouatta palliata* | mantled howler monkey | female | Homogametic | SRR9655168 | SAMN10407156 |
| *Alouatta palliata* | mantled howler monkey | female | Homogametic | SRR9655169 | SAMN10407140 |
| *Anolis carolinensis* | green anole | male | Heterogametic | SRR5508304 | SAMN06891216 |
| *Anolis carolinensis* | green anole | female | Homogametic | SRR5508303 | SAMN06891217 |
| *Calidris pugnax* | ruff | faeder/male | Heterozygotic inversion | ERR1001620 | SAMEA3522198 |
| *Calidris pugnax* | ruff | faeder/male | Heterozygotic inversion | ERR1001621 | SAMEA3522198 |
| *Calidris pugnax* | ruff | resident/male | Homozygotic inversion | ERR1001618 | SAMEA3522199 |
| *Drosophila miranda* | fruit fly | male | Heterogametic | SRR1738163 | SAMN03220545 |
| *Drosophila miranda* | fruit fly | female | Homogametic | SRR1738162 | SAMN03220544 |
| *Nothobranchius furzeri* | turquoise killifish | male | Heterogametic | ERR583467 | SAMEA2698541 |
| *Nothobranchius furzeri* | turquoise killifish | male | Heterogametic | ERR583468 | SAMEA2698543 |
| *Nothobranchius furzeri* | turquoise killifish | female | Homogametic | ERR583470 | SAMEA2698544 |
| *Ornithorhynchus anatinus* | platypus | male | Heterogametic | SRR648711 | SAMN01886049 |
| *Ornithorhynchus anatinus* | platypus | female | Homogametic | SRR648716 | SAMN01886050 |
| *Poecilia reticulata* | guppy | male | Heterogametic | ERR1016729 | SAMEA3539941 |
| *Poecilia reticulata* | guppy | male | Heterogametic | ERR1016727 | SAMEA3539940 |
| *Poecilia reticulata* | guppy | male | Heterogametic | ERR1016717 | SAMEA3539935 |
| *Poecilia reticulata* | guppy | male | Heterogametic | ERR1016716 | SAMEA3539935 |
| *Poecilia reticulata* | guppy | male | Heterogametic | ERR1016726 | SAMEA3539940 |
| *Poecilia reticulata* | guppy | male | Heterogametic | ERR1016728 | SAMEA3539941 |
| *Poecilia reticulata* | guppy | male | Heterogametic | ERR1016730 | SAMEA3539941 |
| *Poecilia reticulata* | guppy | female | Homogametic | ERR1016723 | SAMEA3539938 |
| *Poecilia reticulata* | guppy | female | Homogametic | ERR1016721 | SAMEA3539937 |
| *Poecilia reticulata* | guppy | female | Homogametic | ERR1016725 | SAMEA3539939 |
| *Poecilia reticulata* | guppy | female | Homogametic | ERR1016720 | SAMEA3539937 |
| *Poecilia reticulata* | guppy | female | Homogametic | ERR1016722 | SAMEA3539938 |
| *Poecilia reticulata* | guppy | female | Homogametic | ERR1016724 | SAMEA3539939 |
| *Poecilia reticulata* | guppy | female | Homogametic | ERR1016731 | SAMEA3539942 |
| *Poecilia reticulata* | guppy | female | Homogametic | ERR1016732 | SAMEA3539942 |
| *Poecilia reticulata* | guppy | female | Homogametic | ERR1016733 | SAMEA3539943 |
| *Poecilia reticulata* | guppy | female | Homogametic | ERR1016737 | SAMEA3539945 |
| *Poecilia reticulata* | guppy | female | Homogametic | ERR1016735 | SAMEA3539944 |
| *Poecilia reticulata* | guppy | female | Homogametic | ERR1016738 | SAMEA3539945 |
| *Poecilia reticulata* | guppy | female | Homogametic | ERR1016719 | SAMEA3539936 |
| *Poecilia reticulata* | guppy | female | Homogametic | ERR1016718 | SAMEA3539936 |
| *Poecilia reticulata* | guppy | female | Homogametic | ERR1016734 | SAMEA3539943 |
| *Poecilia reticulata* | guppy | female | Homogametic | ERR1016736 | SAMEA3539944 |
| *Pogona vitticeps* | central bearded dragon | ZW female | Heterogametic | ERR409918 | SAMEA2300449 |
| *Pogona vitticeps* | central bearded dragon | ZW female | Heterogametic | ERR409919 | SAMEA2300449 |
| *Pogona vitticeps* | central bearded dragon | ZW female | Heterogametic | ERR409920 | SAMEA2300449 |
| *Pogona vitticeps* | central bearded dragon | ZZ male | Homogametic | ERR409943 | SAMEA2300447 |
| *Pogona vitticeps* | central bearded dragon | ZZ male | Homogametic | ERR409944 | SAMEA2300447 |
| *Pogona vitticeps* | central bearded dragon | ZZ male | Homogametic | ERR409945 | SAMEA2300447 |

Supplementary Table 2: Reference genomes used as either study-species or synteny-species reference genomes (see Figure 2a in Main text). Only reference genomes from homogametic samples were used as study-species reference genomes. For the two heterogametic genomes (*G. gallus* and *H. sapiens*), the sex-limited chromosome was removed prior to analyses.

| Species | Common name | Assembly ID | GenBank accession | Sex | Sex chromosomes |
|---|---|---|---|---|---|
| *Alauda arvensis* | Eurasian skylark | skylark_min1kb | NA* | male | ZZ (Homogametic) |
| *Alouatta palliata* | mantled howler monkey | AloPal_v1_BIUU | GCA_004027835.1 | female | XX (Homogametic) |
| *Anolis carolinensis* | green anole | AnoCar2.0 | GCA_000090745.2 | female | XX (Homogametic) |
| *Calidris pugnax* | ruff | ASM143184v1 | GCA_001431845.1 | not collected | NA |
| *Drosophila miranda* | fruit fly | DroMir_2.2 | GCA_000269505.2 | female | XX (Homogametic) |
| *Gallus gallus* | chicken | GRCg6a | GCA_000002315.5 | female | ZW (Heterogametic) |
| *Homo sapiens* | human | GRCh38.p13 | GCA_000001405.28 | male | XY (Heterogametic) |
| *Nothobranchius furzeri* | turquoise killifish | Nfu_20140520 | GCA_001465895.2 | female | XX (Homogametic) |
| *Ornithorhynchus anatinus* | platypus | ASM227v2/ornAna2 | GCA_000002275.2 | female | XX (Homogametic) |
| *Poecilia reticulata* | guppy | Guppy_female_1.0+MT | GCA_000633615.2 | female | XX (Homogametic) |
| *Pogona vitticeps* | central bearded dragon | pvi1.1 | GCA_900067755.1 | male | ZZ (Homogametic) |
| *Suricata suricatta* | meerkat | meerkat_22Aug2017_6uvM2_HiC | GCA_006229205.1 | female | XX (Homogametic) |
| *Taeniopygia guttata* | zebra finch | taeGut3.2.4 | GCA_000151805.2 | male | ZZ (Homogametic) |

* Available at `https://doi.org/10.5061/dryad.95x69p8f9`

Supplementary Table 3: Information on what sex chromosome(s)/scaffold(s) were detected with the pipeline, and the findZX configuration and unit files used to run each analysis. See also Supplementary Methods for more details on sex chromosome systems of each species, and commands to run each analysis.

| | ***See Supplementary Table 2 for details on reference genomes*** | | | ***Files found on GitHub in directory: config/9_species_config/*** | | |
|---|---|---|---|---|---|---|
| **Study-species** | **Study-species reference genome** | **Synteny-species reference genome** | **Program** | **Config file** | **Unit file** | **# individuals** |
| *A. arvensis* | *A. arvensis* | NA | findZX (-R modify_genome) | Alauda_arvensis_config.yml | Alauda_arvensis_units.tsv | 2 |
| *A. arvensis* | *A. arvensis (consensus)* | *T. guttata (Z;3;4A;5*)* | findZX-synteny | Alauda_arvensis_consensus_config.yml | Alauda_arvensis_units.tsv | 2 |
| *A. arvensis* | *A. arvensis* | *G. gallus (Z;3;4;5*)* | findZX-synteny | Alauda_arvensis_consensus_config.yml | Alauda_arvensis_units.tsv | 2 |
| *A. palliata* | *A. palliata* | NA | findZX | Alouatta_palliata_config.yml | Alouatta_palliata_units.tsv | 4 |
| *A. palliata* | *A. palliata* | *H. sapiens (X;3*)* | findZX-synteny | Alouatta_palliata_config.yml | Alouatta_palliata_units.tsv | 4 |
| *A. palliata* | *A. palliata* | *H. sapiens (X;3*)* | findZX-synteny | Alouatta_palliata_config_1M_1F.yml | Alouatta_palliata_units_1M_1F.tsv | 2 |
| *A. palliata* | *A. palliata* | *H. sapiens (X;3*)* | findZX-synteny | Alouatta_palliata_subsample_1M_1F_config.yml | Alouatta_palliata_units_1M_1F.tsv | 2 |
| *A. palliata* | *A. palliata* | *S. suricatta (X;5*)* | findZX-synteny | Alouatta_palliata_config.yml | Alouatta_palliata_units.tsv | 4 |
| *A. carolinensis* | *A. carolinensis (LGb*)* | *NA* | findZX | Anolis_carolinensis_config.yml | Anolis_carolinensis_units.tsv | 2 |
| *C. pugnax* | *C. pugnax (scaffold28/KQ482164.1*)* | *NA* | findZX | Calidris_pugnax_config.yml | Calidris_pugnax_units.tsv | 3 |
| *C. pugnax* | *C. pugnax* | *G. gallus (11*)* | findZX-synteny | Calidris_pugnax_config.yml | Calidris_pugnax_units.tsv | 3 |
| *D. miranda* | *D. miranda (XL;XR;3*)* | *NA* | findZX | Drosophila_miranda_config.yml | Drosophila_miranda_units.tsv | 2 |
| *N. furzeri* | *N. furzeri (sgr05*)* | *NA* | findZX | Nothobranchius_furzeri_config.yml | Nothobranchius_furzeri_units.tsv | 3 |
| *N. furzeri* | *N. furzeri (sgr05*)* | *NA* | findZX | Nothobranchius_furzeri_config_1M_1F.yml | Nothobranchius_furzeri_units_1M_1F.tsv | 2 |
| *O. anatinus* | *O. anatinus (X1;X2;X3;X4;X5*)* | *NA* | findZX | Ornithorhynchus_anatinus_config.yml | Ornithorhynchus_anatinus_units.tsv | 2 |
| *P. reticulata* | *P. reticulata (LG12*)* | *NA* | findZX | Poecilia_reticulata_config.yml | Poecilia_reticulata_units.tsv | 23 |
| *P. reticulata* | *P. reticulata (LG12*)* | *NA* | findZX | Poecilia_reticulata_config_1M_1F.yml | Poecilia_reticulata_units_1M_1F.tsv | 2 |
| *P. vitticeps* | *P. vitticeps* | *G. gallus (17;23**)* | findZX | Pogona_vitticeps_config.yml | Pogona_vitticeps_units.tsv | 2(6)*** |

* Sought after chromosomes/scaffolds identified with the pipeline
** Sought after chromosomes/scaffolds not identified with the pipeline
*** 2 individuals, each sequenced three times (see Supplementary Table 2)

Supplementary Table 4: Average genome coverage per studied sample. Genome coverage values are given for all three mismatch settings ("unfiltered", "≤2 mismatches", and "0 mismatches").

| **Study species / Reference genome** | **Sample** | **Homogametic/Heterogametic** | **unfiltered** | **≤2 mismatches** | **0 mismatches** |
|---|---|---|---|---|---|
| *Alauda arvensis* | SRR10340221 | Heterogametic | 29.54 | 14.12 | 4.76 |
| *Alauda arvensis* | SRR10340220 | Homogametic | 29.58 | 24.09 | 17.44 |
| *Alauda arvensis (consensus genome)* | SRR10340221 | Heterogametic | 26.15 | 17.46 | 7.99 |
| *Alauda arvensis (consensus genome)* | SRR10340220 | Homogametic | 27.18 | 19.91 | 8.69 |
| *Alouatta palliata* | SRR9655170 | Heterogametic | 3.05 | 2.88 | 2.38 |
| *Alouatta palliata* | SRR9655170 (subset) | Heterogametic | 1.74 | 1.69 | 1.43 |
| *Alouatta palliata* | SRR9655171 | Heterogametic | 5.11 | 4.86 | 4.04 |
| *Alouatta palliata* | SRR9655168 | Homogametic | 5.04 | 4.82 | 4.01 |
| *Alouatta palliata* | SRR9655168 (subset) | Homogametic | 1.97 | 1.91 | 1.63 |
| *Alouatta palliata* | SRR9655169 | Homogametic | 6.86 | 6.58 | 5.49 |
| *Anolis carolinensis* | SRR5508304 | Heterogametic | 13.64 | 12.48 | 8.38 |
| *Anolis carolinensis* | SRR5508303 | Homogametic | 15.78 | 14.53 | 10.56 |
| *Calidris pugnax* | ERR1001620 | Heterogametic | 7.29 | 6.47 | 4.75 |
| *Calidris pugnax* | ERR1001621 | Heterogametic | 7.92 | 7.01 | 5.19 |
| *Calidris pugnax* | ERR1001618 | Homogametic | 8.31 | 7.39 | 5.50 |
| *Drosophila miranda* | SRR1738163 | Heterogametic | 13.94 | 11.33 | 8.16 |
| *Drosophila miranda* | SRR1738162 | Homogametic | 8.02 | 6.77 | 5.34 |
| *Nothobranchius furzeri* | ERR583467 | Heterogametic | 19.64 | 17.41 | 13.47 |
| *Nothobranchius furzeri* | ERR583468 | Heterogametic | 30.94 | 27.75 | 21.53 |
| *Nothobranchius furzeri* | ERR583470 | Homogametic | 29.24 | 26.48 | 22.52 |
| *Ornithorhynchus anatinus* | SRR648711 | Heterogametic | 12.24 | 11.39 | 6.74 |
| *Ornithorhynchus anatinus* | SRR648716 | Homogametic | 12.83 | 12.16 | 9.11 |
| *Poecilia reticulata* | ERR1016716 | Heterogametic | 3.20 | 2.84 | 2.18 |
| *Poecilia reticulata* | ERR1016717 | Heterogametic | 6.83 | 6.09 | 4.65 |
| *Poecilia reticulata* | ERR1016726 | Heterogametic | 3.00 | 2.66 | 1.99 |
| *Poecilia reticulata* | ERR1016727 | Heterogametic | 3.45 | 3.05 | 2.29 |
| *Poecilia reticulata* | ERR1016728 | Heterogametic | 3.72 | 3.29 | 2.48 |
| *Poecilia reticulata* | ERR1016729 | Heterogametic | 6.19 | 5.48 | 4.09 |
| *Poecilia reticulata* | ERR1016730 | Heterogametic | 7.87 | 6.97 | 5.26 |
| *Poecilia reticulata* | ERR1016718 | Homogametic | 5.09 | 4.52 | 3.39 |
| *Poecilia reticulata* | ERR1016719 | Homogametic | 10.73 | 9.54 | 7.15 |
| *Poecilia reticulata* | ERR1016720 | Homogametic | 3.77 | 3.34 | 2.47 |
| *Poecilia reticulata* | ERR1016721 | Homogametic | 4.35 | 3.85 | 2.86 |
| *Poecilia reticulata* | ERR1016722 | Homogametic | 2.57 | 2.26 | 1.66 |
| *Poecilia reticulata* | ERR1016723 | Homogametic | 2.93 | 2.58 | 1.90 |
| *Poecilia reticulata* | ERR1016724 | Homogametic | 7.93 | 7.07 | 5.29 |
| *Poecilia reticulata* | ERR1016725 | Homogametic | 9.11 | 8.12 | 6.09 |
| *Poecilia reticulata* | ERR1016731 | Homogametic | 3.51 | 3.12 | 2.34 |
| *Poecilia reticulata* | ERR1016732 | Homogametic | 7.34 | 6.52 | 4.89 |
| *Poecilia reticulata* | ERR1016733 | Homogametic | 4.22 | 3.73 | 2.76 |
| *Poecilia reticulata* | ERR1016734 | Homogametic | 4.78 | 4.23 | 3.14 |
| *Poecilia reticulata* | ERR1016735 | Homogametic | 3.05 | 2.71 | 2.03 |
| *Poecilia reticulata* | ERR1016736 | Homogametic | 6.39 | 5.69 | 4.26 |
| *Poecilia reticulata* | ERR1016737 | Homogametic | 8.29 | 7.36 | 5.43 |
| *Poecilia reticulata* | ERR1016738 | Homogametic | 9.47 | 8.41 | 6.24 |
| *Pogona vitticeps* | ERR409918 | Heterogametic | 8.23 | 6.24 | 2.78 |
| *Pogona vitticeps* | ERR409919 | Heterogametic | 12.62 | 7.75 | 2.91 |
| *Pogona vitticeps* | ERR409920 | Heterogametic | 12.43 | 7.60 | 2.84 |
| *Pogona vitticeps* | ERR409943 | Homogametic | 13.67 | 11.24 | 6.72 |
| *Pogona vitticeps* | ERR409944 | Homogametic | 12.98 | 10.01 | 5.50 |
| *Pogona vitticeps* | ERR409945 | Homogametic | 7.14 | 5.58 | 2.96 |

### 3 Supplementary Figures

#### 3.1 Example output - plot types 1-5

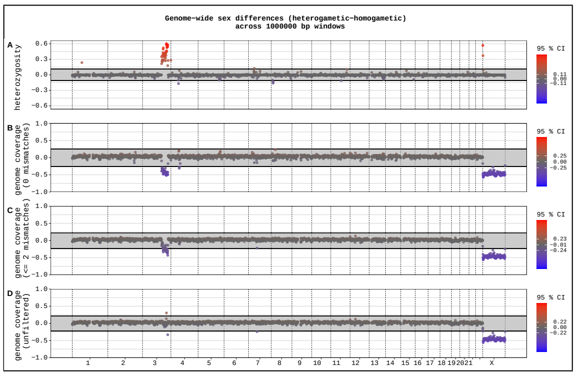


(**A**) Plot type 1

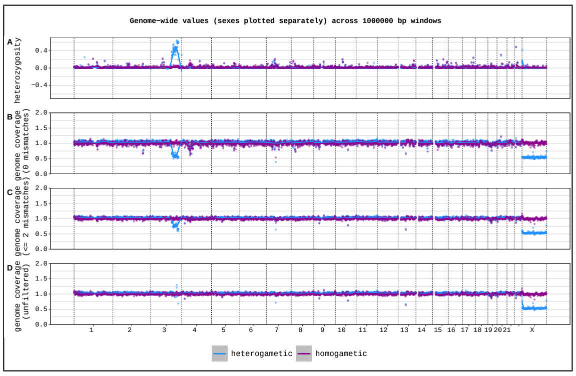


(**B**) Plot type 2

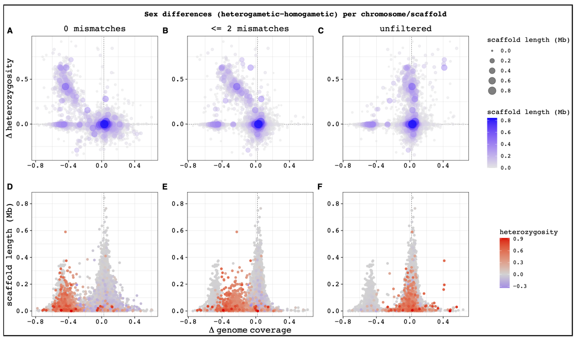


(**C**) Plot type 3

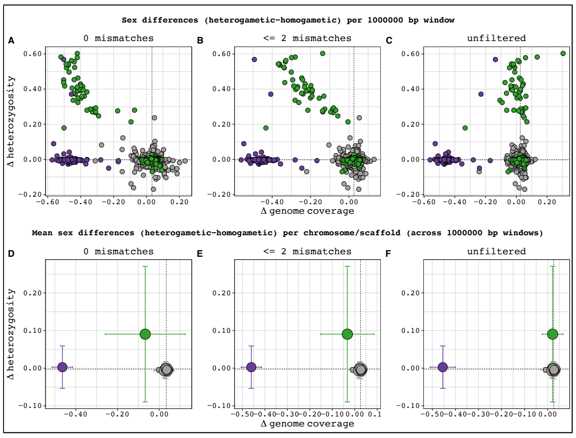


(**D**) Plot type 4

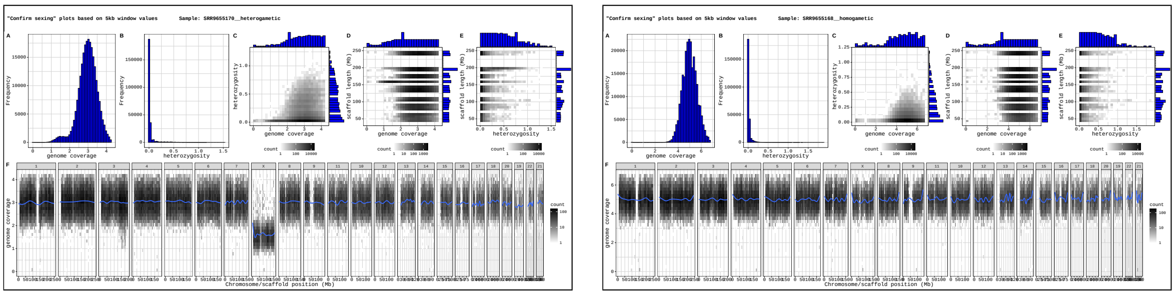


(**E**) Plot type 5 (two different individuals)

Supplementary Figure 1: FindZX will produce five types of output plots (here shown as miniatures **A-E)**, all based on heterozygosity and genome coverage values. **(A):** Genome-wide sex differences across specified genome windows (here 1 Mb; see Figure 3 in main text). **(B):** Genome-wide sex-specific values across specified genome windows (here 1 Mb; see Supplementary Figure 2). **(C):** Per chromosome/scaffold values (bottom row also include chromosome/scaffold length; see Figure 4 in main text). **(D):** Genome window values (here 1 Mb; upper row), and per-chromosome/scaffold values based on the same windows (bottom row; see Figure 5 in main text). **(E):** "Confirm sexing" plots. Differences in genome coverage and heterozygosity profiles may reveal if some samples were given the wrong sex (see Supplementary Figure 3 and 4).

#### 3.2 Example output - plot type 2

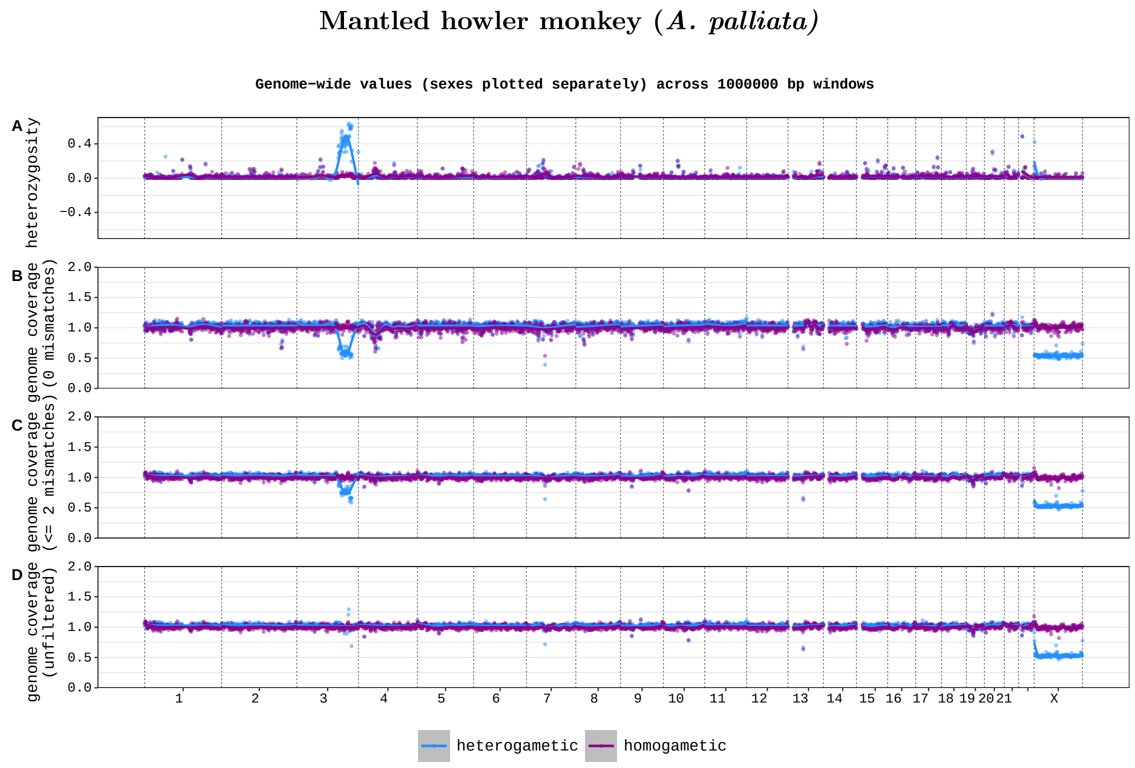


Supplementary Figure 2: Per-sex genome coverage and heterozygosity values (1 Mb windows) for mantled howler monkey, plotted along chromosome positions in the human genome. The four rows show: (**A**) heterozygosity, and genome coverage with (**B**) strict filtering (0 mismatches allowed), (**C**) intermediate filtering ($\leq$2 mismatches) and (**D**) no filtering of mapped reads (unfiltered). The values for each sex are plotted separately, with a smoothing line for each sex (heterogametic in blue, homogametic in purple). The data confirms that chromosome X and a part of chromosome 3 are sex-linked in this species [4].

#### 3.3 Example output - plot type 5

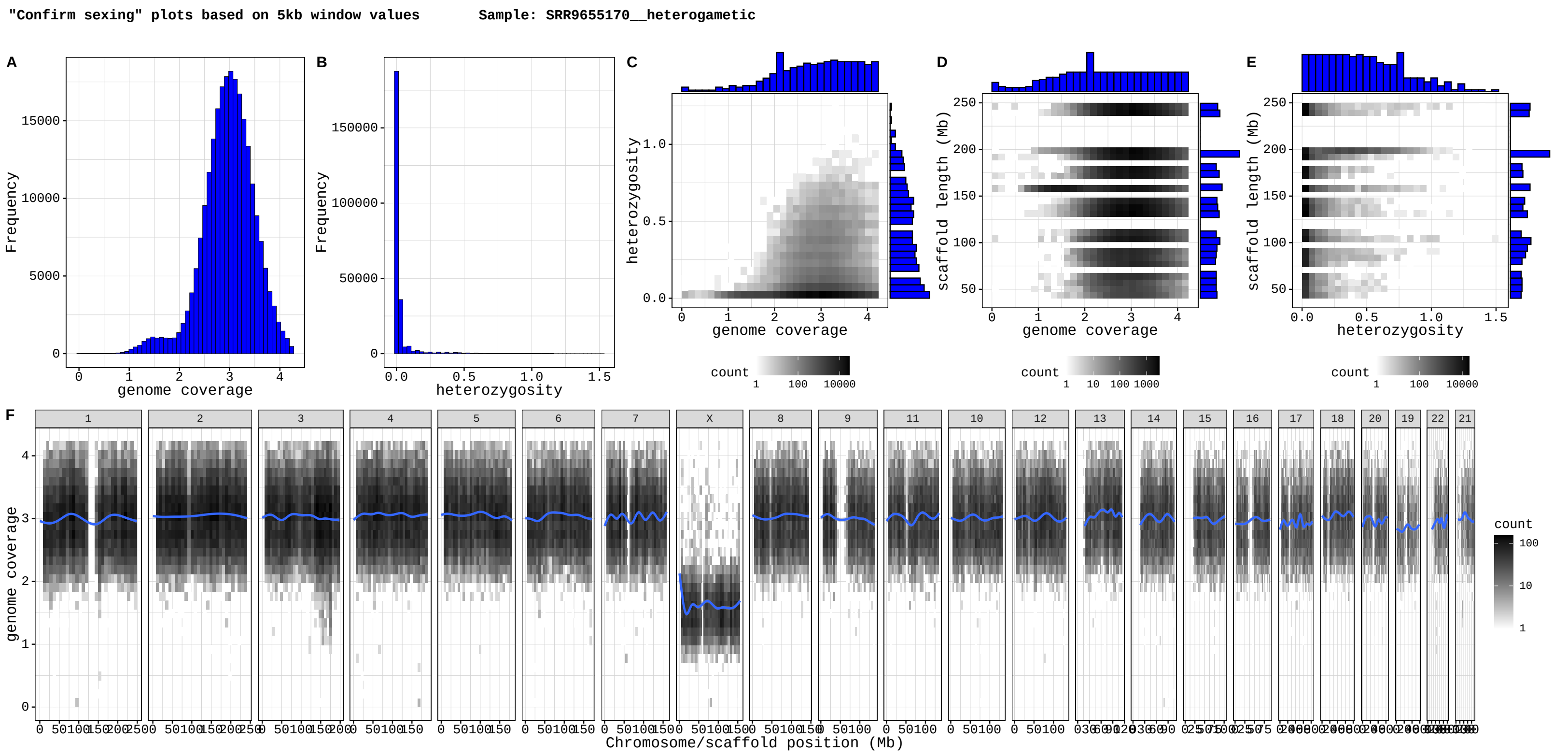


Supplementary Figure 3: Example of a "Confirm-sexing" plot (see Figure 2 in main text), from a male (XY) mantled howler monkey sample (*A. palliata*).

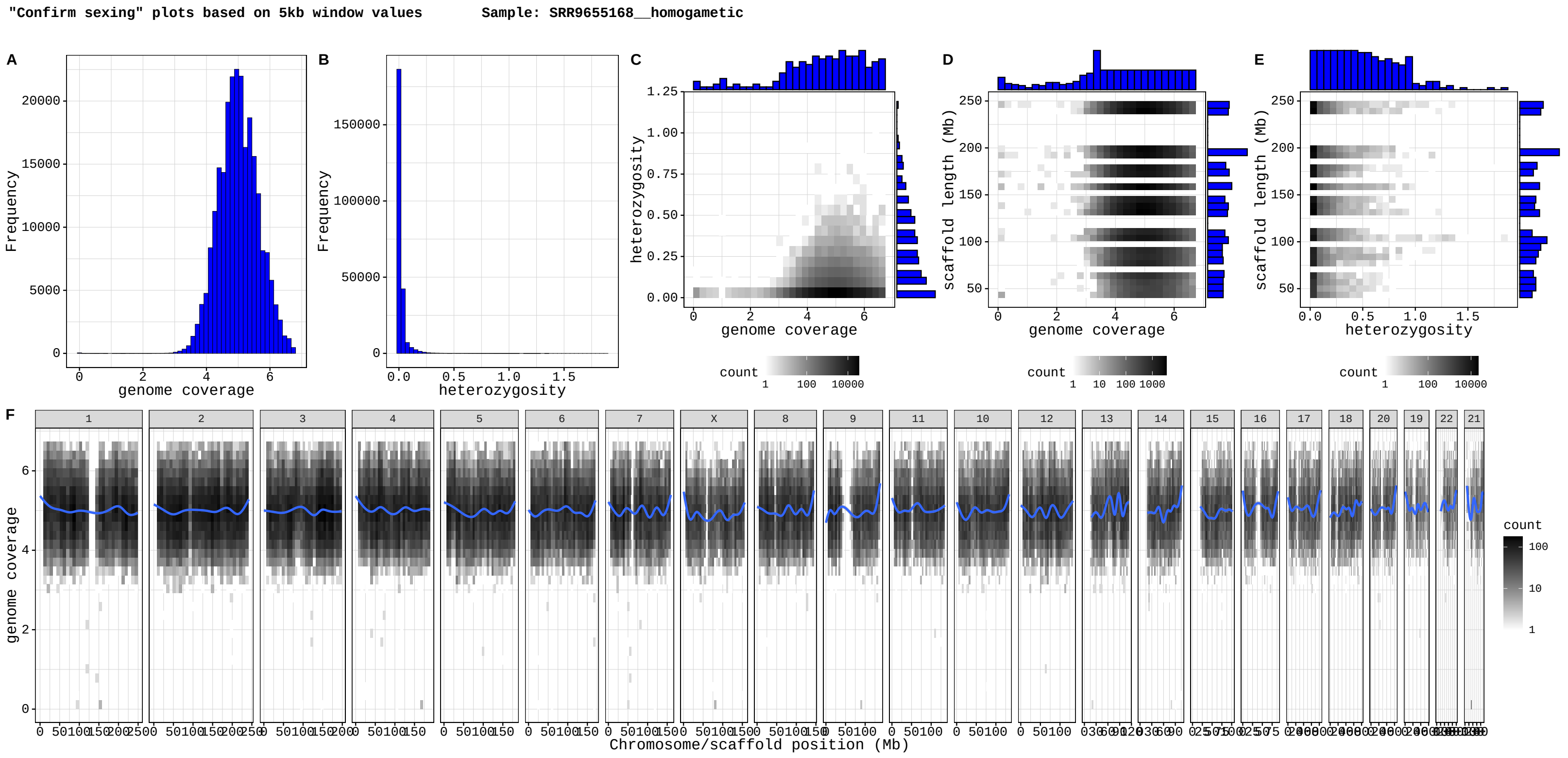


Supplementary Figure 4: Example of a "Confirm-sexing" plot (see Figure 2 in main text), from a female (XX) mantled howler monkey sample (*A. palliata*).

#### 3.4 Output plots type 4 from all nine studied species

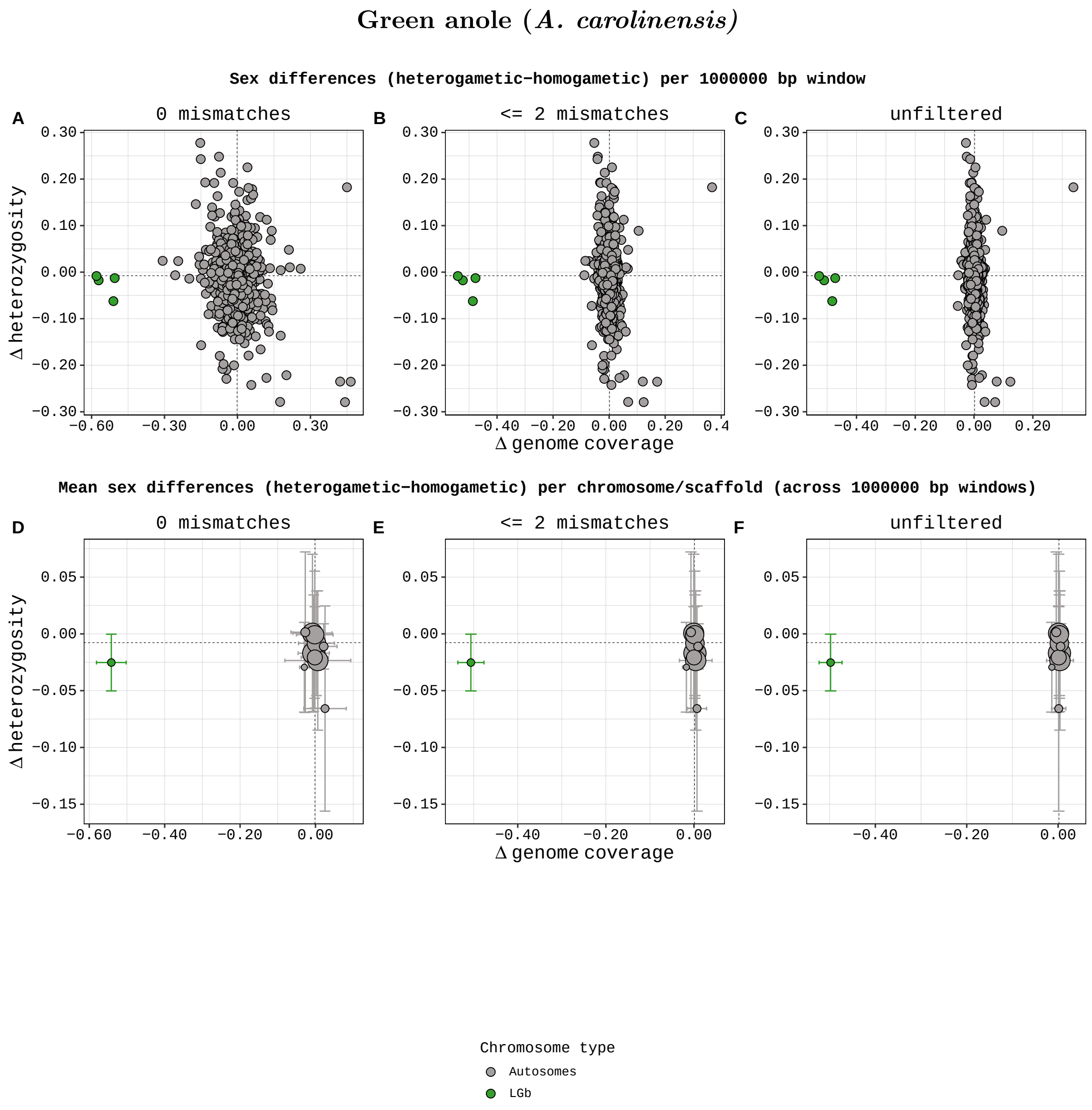


Supplementary Figure 5: **A-C)** Sex differences in genome coverage and heterozygosity for all 1 Mb genome windows. **D-F)** Mean (± SD) sex differences in genome coverage and heterozygosity per chromosome/scaffold, calculated from the 1 Mb genome windows. Dashed lines mark the genome-wide median across all 1 Mb windows. Data from 1 male and 1 female green anole (*A. carolinensis*), analysed using findZX (without the use of a synteny-species reference genome). The previously identified sex chromosome in this species (LGb [5]) is a clear outlier.

#### Platypus (*O. anatinus*)

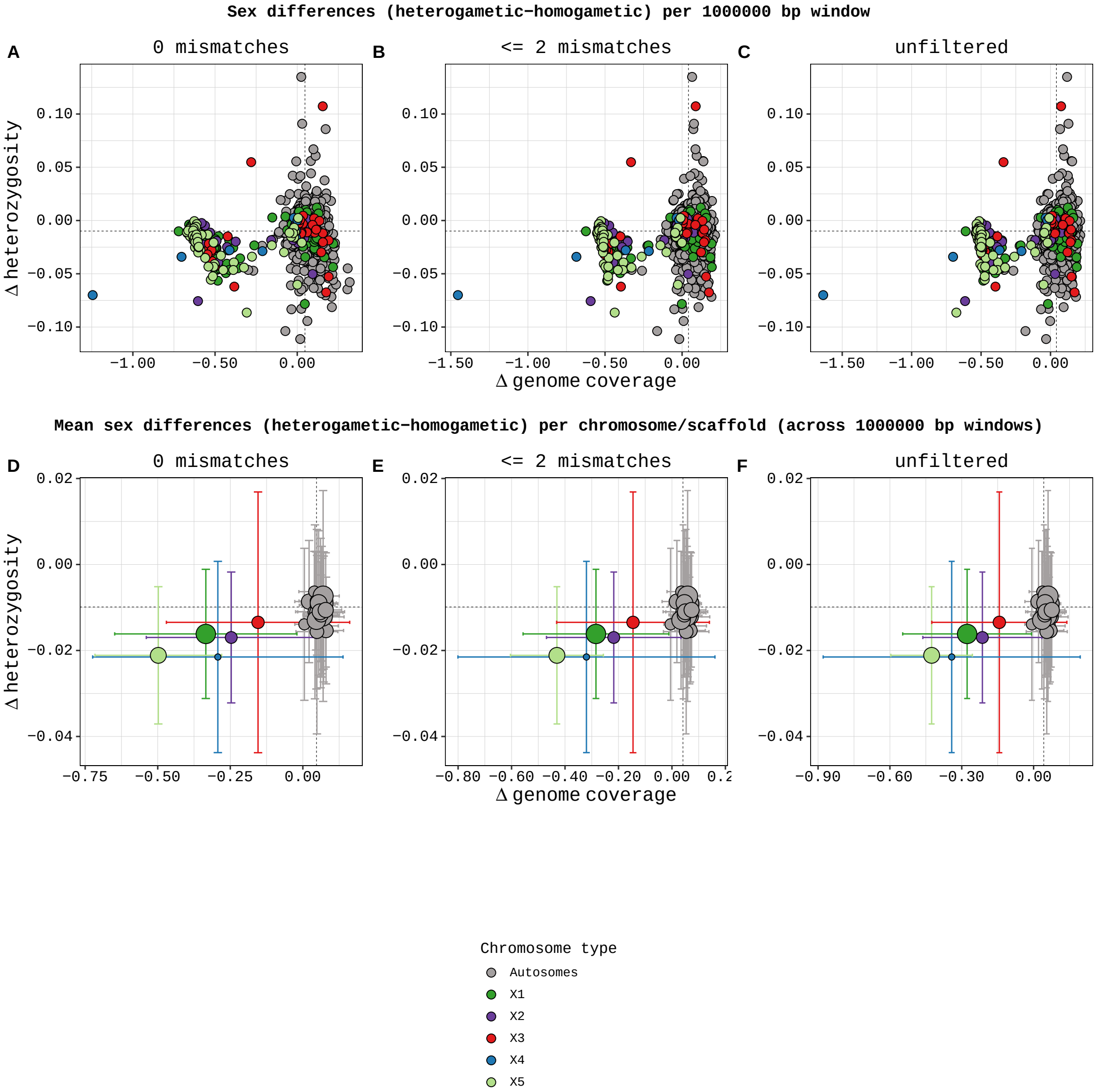


Supplementary Figure 6: **A-C)** Sex differences in genome coverage and heterozygosity for all 1 Mb genome windows. **D-F)** Mean (± SD) sex differences in genome coverage and heterozygosity per chromosome/scaffold, calculated from the 1 Mb genome windows. Dashed lines mark the genome-wide median across all 1 Mb windows. Data from 1 male and 1 female platypus (*O. anatinus*), analysed using findZX (without the use of a synteny-species reference genome. The previously identified sex chromosomes in this species ($X_1$, $X_2$, $X_3$, $X_4$ and $X_5$ [9]) are clear outliers.

#### Mantled howler monkey (*A. palliata*)

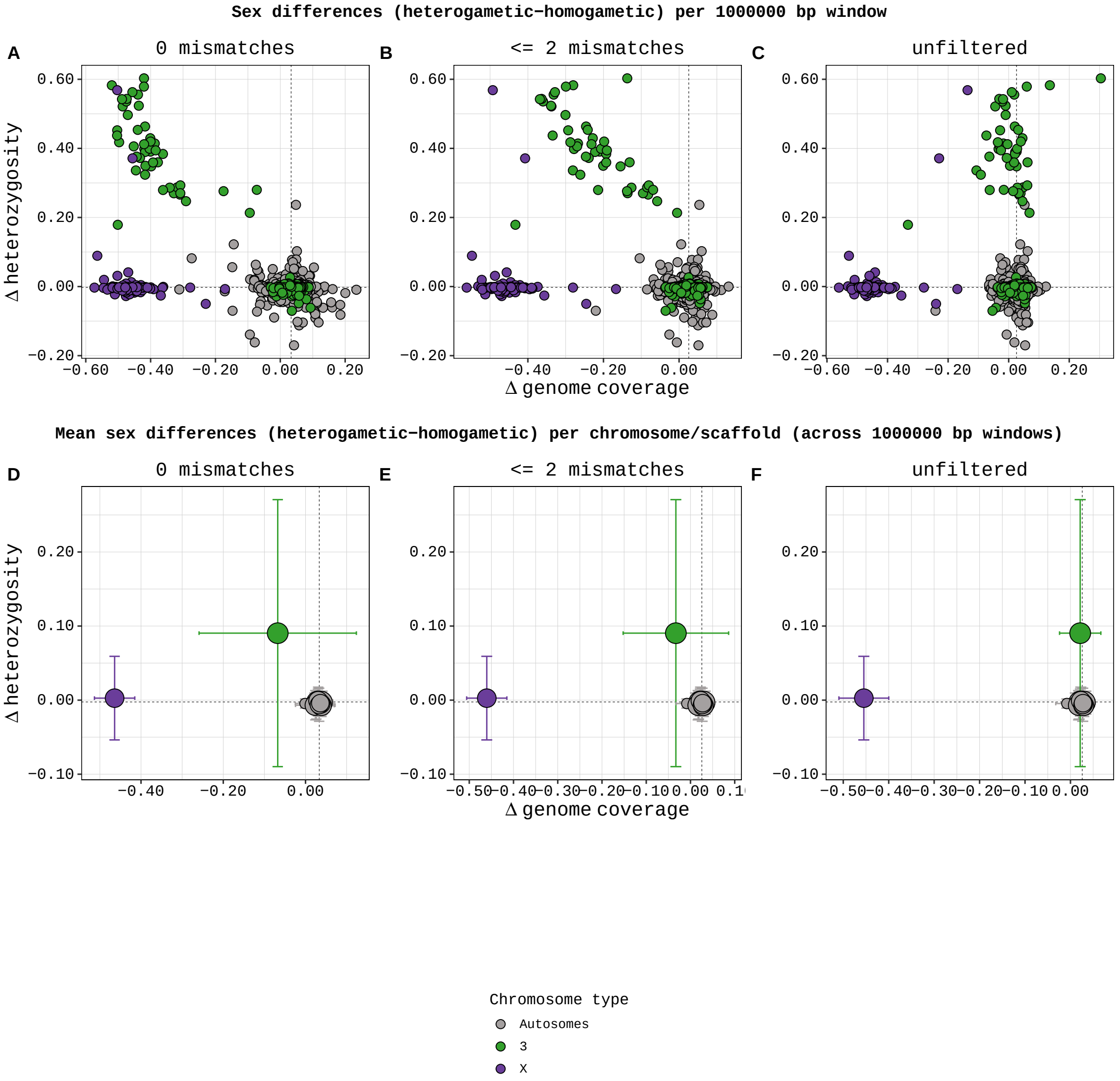


Supplementary Figure 7: **A-C)** Sex differences in genome coverage and heterozygosity for all 1 Mb genome window. **D-F)** Mean (± SD) sex differences in genome coverage and heterozygosity per chromosome/scaffold, calculated from the 1 Mb genome windows. Dashed lines mark the genome-wide median across all 1 Mb windows. Data from 2 male and 2 female mantled howler monkeys (*A. palliata*), analysed using findZX-synteny and human (*H. sapiens*) as a synteny-species reference genome. The previously identified sex chromosomes in this species (X and 3 [4]) are clear outliers.

#### Eurasian skylark (*A. alauda*)

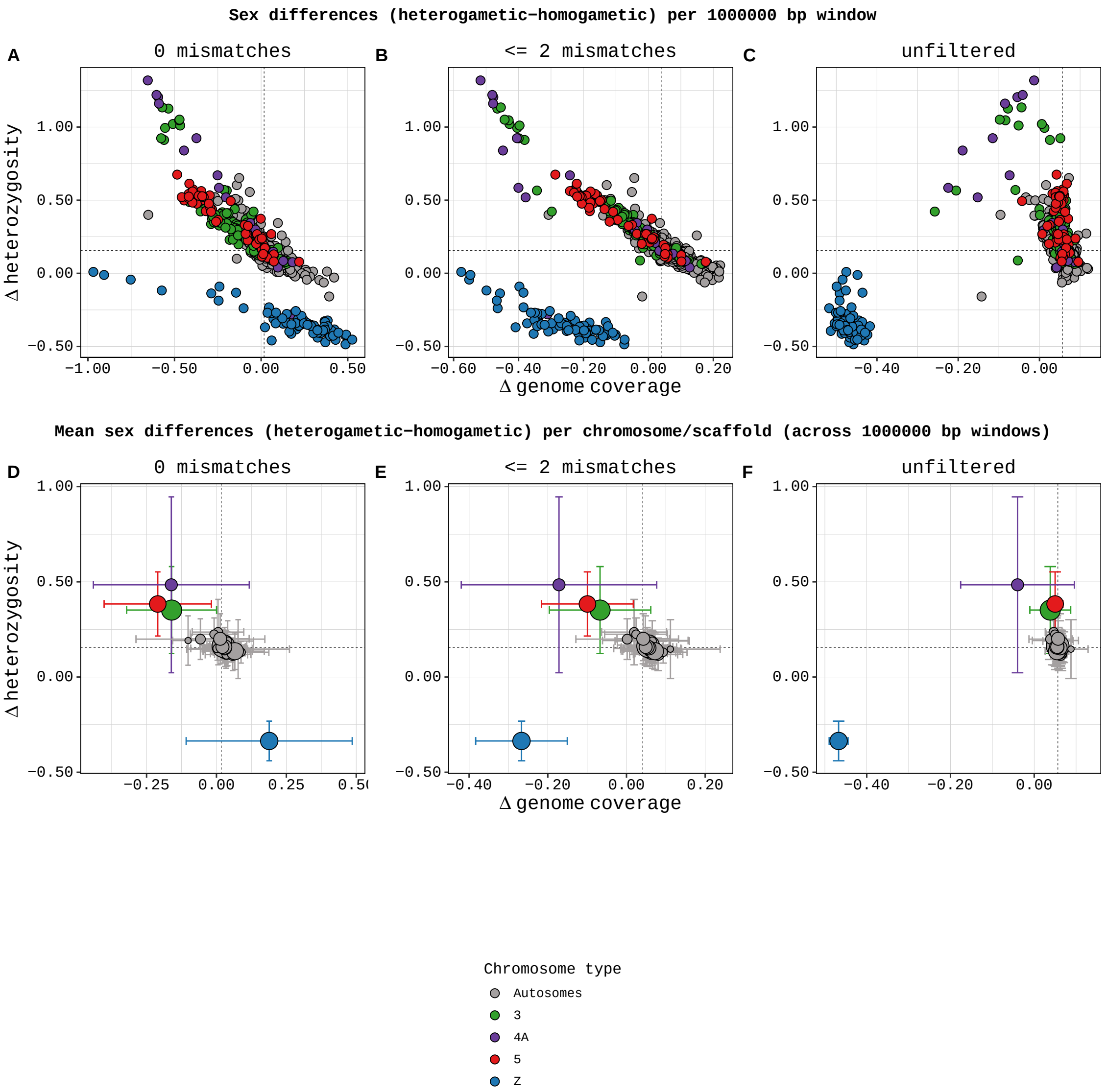


Supplementary Figure 8: **A-C)** Sex differences in genome coverage and heterozygosity for all 1 Mb genome windows. **D-F)** Mean (± SD) sex differences in genome coverage and heterozygosity per chromosome/scaffold, calculated from the 1 Mb genome windows. Dashed lines mark the genome-wide median across all 1 Mb windows. Data from 1 male and 1 female Eurasian skylarks (*A. arvensis*), analysed using findZX-synteny and zebra finch (*T. guttata*) as a synteny-species reference genome. The previously identified sex chromosomes in this species (Z, 3, 4A, and 5 [1, 2]) are clear outliers. The analysis was performed using a "consensus reference genome".

#### Fruit fly (*D. miranda*)

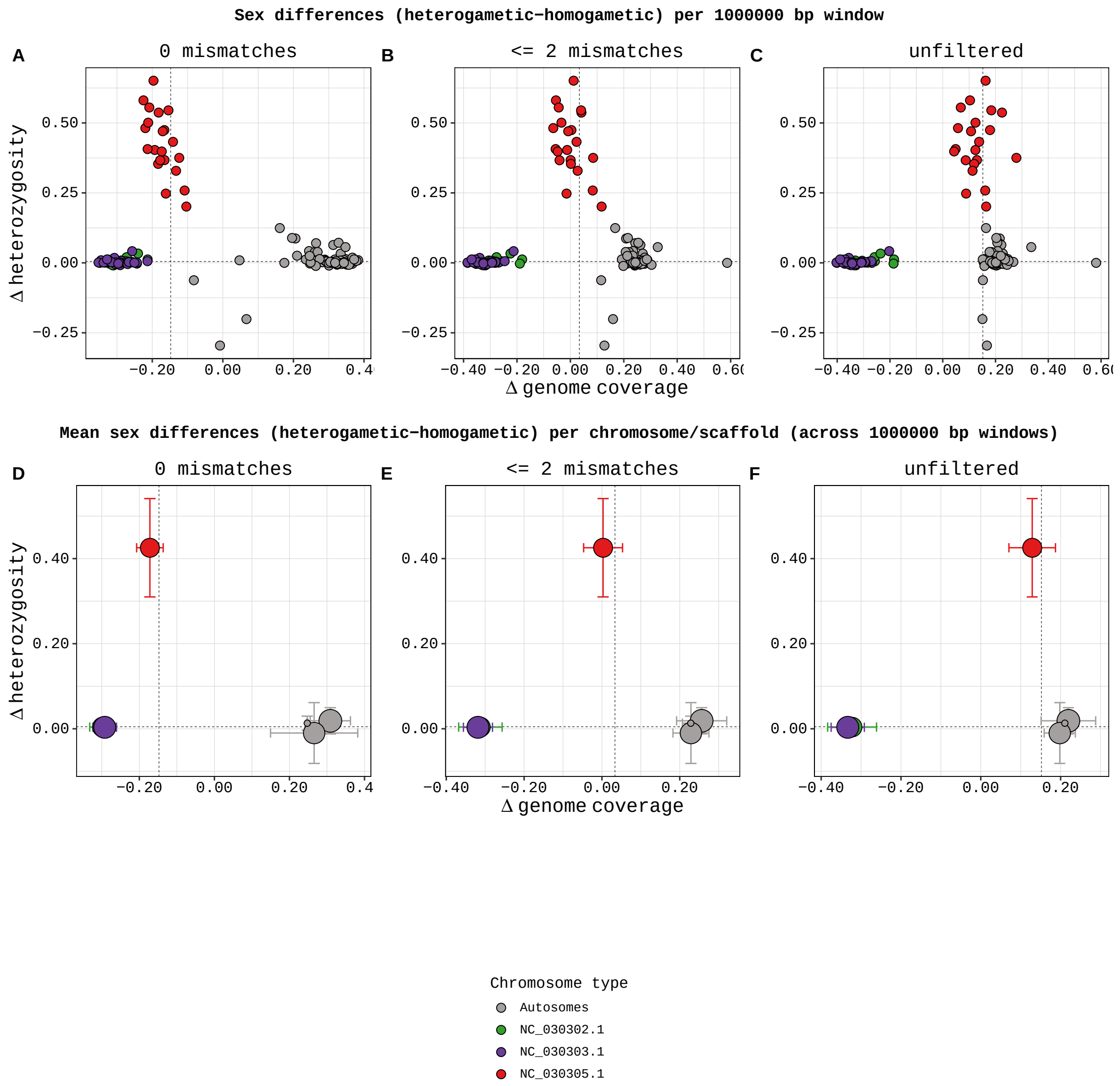


Supplementary Figure 9: **A-C)** Sex differences in genome coverage and heterozygosity for all 1 Mb genome windows. **D-F)** Mean (± SD) sex differences in genome coverage and heterozygosity per chromosome/scaffold, calculated from the 1 Mb genome windows. Dashed lines mark the genome-wide median across all 1 Mb windows. Data from 1 male and 1 female fruit fly (*D. miranda*), analysed using findZX (without the use of a synteny-species reference genome). The previously identified sex chromosomes in this species (XL; NC_030302.1, XR; NC_030303.1 and 3; NC_030305.1 [7]) are clear outliers.

#### Turquoise killifish (*N. furzeri*)

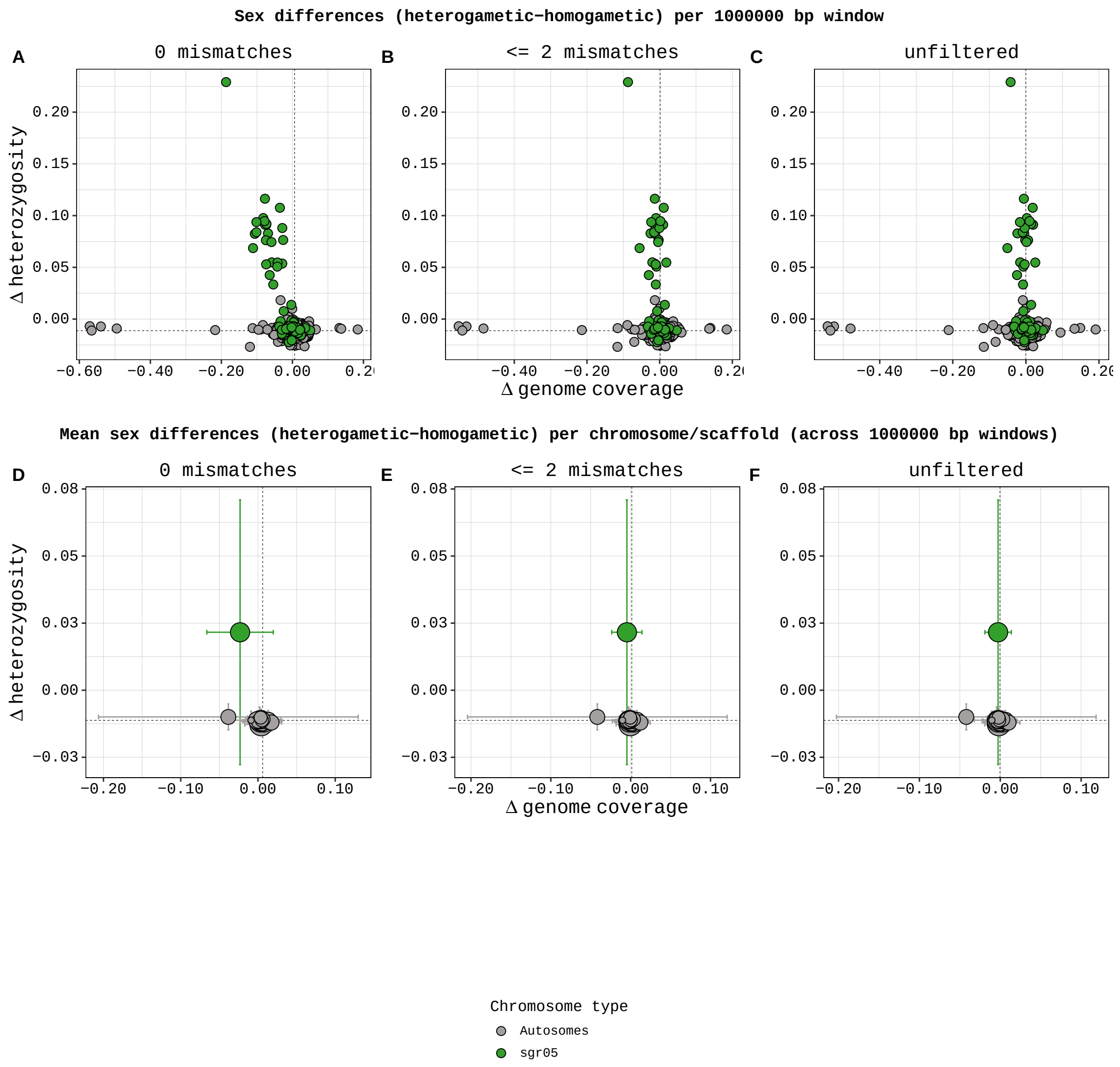


Supplementary Figure 10: **A-C)** Sex differences in genome coverage and heterozygosity for all 1 Mb genome windows. **D-F)** Mean (± SD) sex differences in genome coverage and heterozygosity per chromosome/scaffold, calculated from the 1 Mb genome windows. Dashed lines mark the genome-wide median across all 1 Mb windows. Data from 2 male and 1 female turquoise killifish (*N. furzeri*), analysed using findZX (without a synteny-species reference genome). The previously identified sex chromosome in this species (sgr05 [8]) is a clear outlier.

#### Guppy (*P. reticulata*)

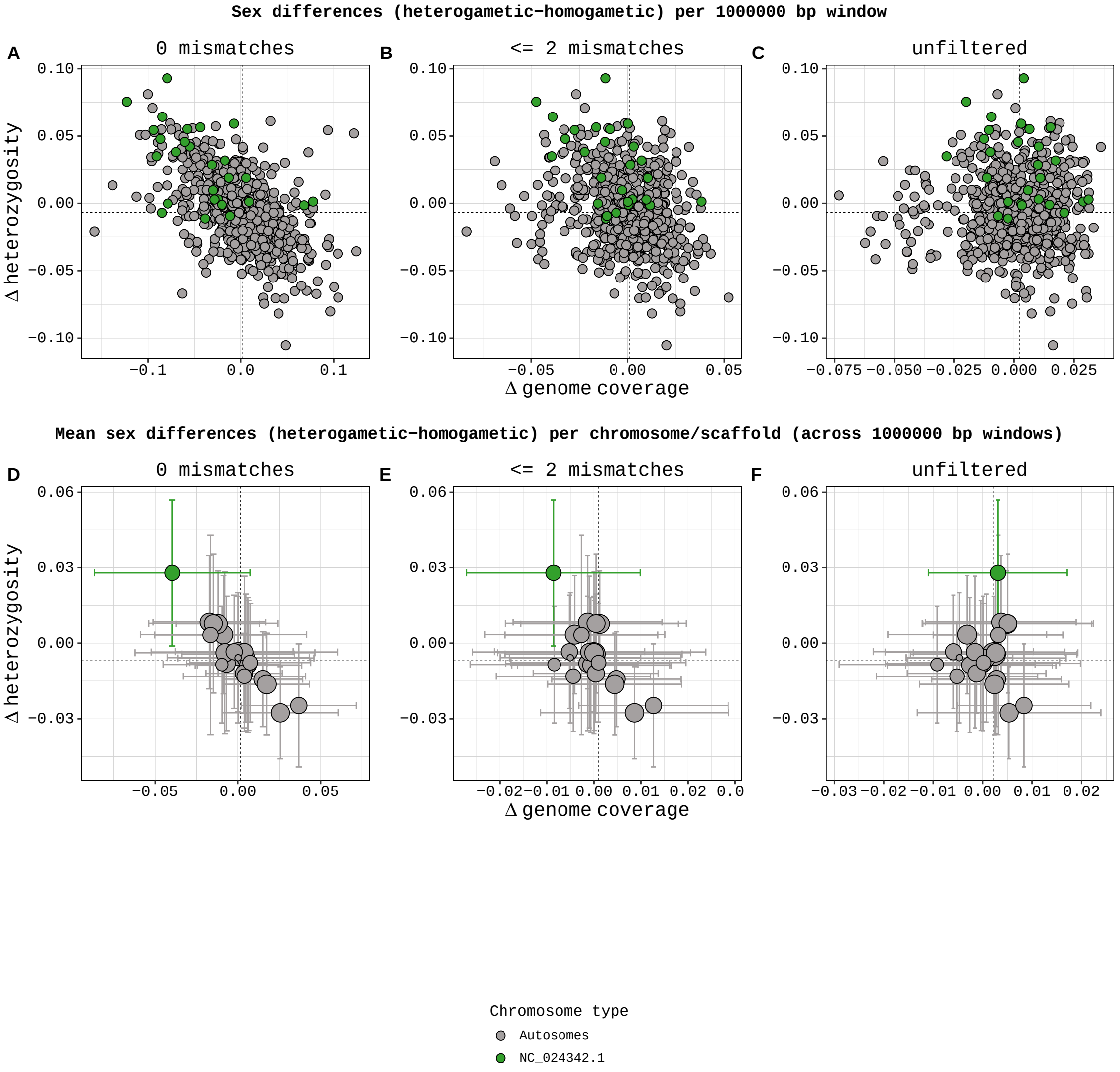


Supplementary Figure 11: **A-C)** Sex differences in genome coverage and heterozygosity for all 1 Mb genome windows. **D-F)** Mean (± SD) sex differences in genome coverage and heterozygosity per chromosome/scaffold, calculated from the 1 Mb genome windows. Dashed lines mark the genome-wide median across all 1 Mb windows. Data from 7 male and 16 female guppies (*P. reticulata*), analysed using findZX (without a synteny-species reference genome). The previously identified sex chromosome in this species (Chromosome 12/NC_024342.1 [10, 11]) is a clear outlier.

### Central bearded dragon (*P. vitticeps*)

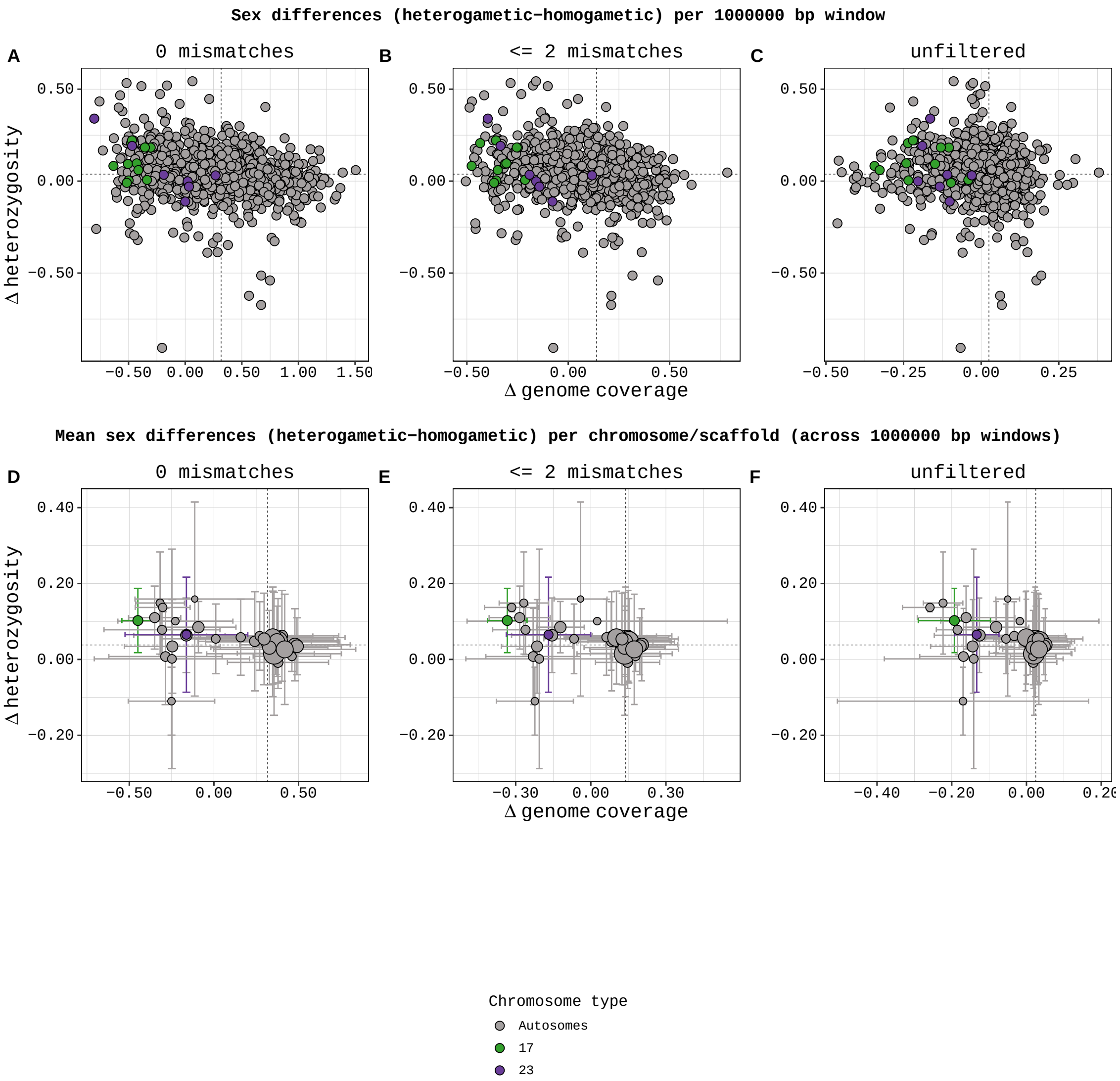


Supplementary Figure 12: **A-C)** Sex differences in genome coverage and heterozygosity for all 1 Mb genome windows. **D-F)** Mean (± SD) sex differences in genome coverage and heterozygosity per chromosome/scaffold, calculated from the 1 Mb genome windows. Dashed lines mark the genome-wide median across all 1 Mb windows. Data from 1 male and 1 female (sequenced three times each; see Supplementary Table 2) central bearded dragons (*P. vitticeps*), analysed using findZX-synteny with the chicken (*G. gallus*) as a synteny-species reference genome. The previously identified sex chromosomes in this species (17 and 23 [12]) are outliers, but not the only ones.

#### Ruff (*C. pugnax*)

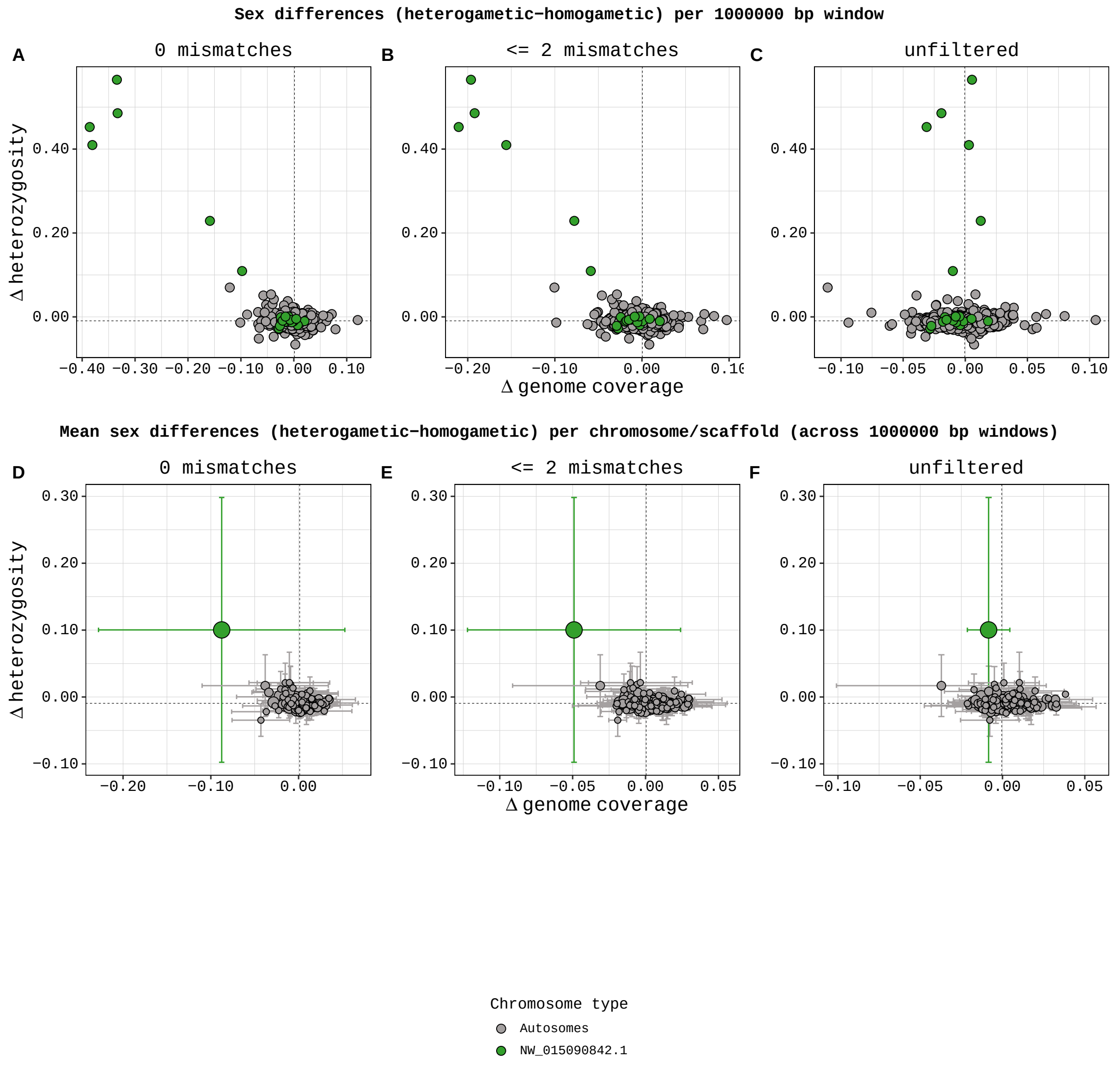


Supplementary Figure 13: **A-C)** Morph differences in genome coverage and heterozygosity for all 1 Mb genome windows. **D-F)** Mean (± SD) morph differences in genome coverage and heterozygosity per chromosome/scaffold, calculated from the 1 Mb genome windows. Dashed lines mark the genome-wide median across all 1 Mb windows. Data from 3 male ruffs (*C. pugnax*); 2 faeder males (heterozygotic inversion) and 1 resident male (homozygotic inversion). The data was analysed using findZX (without the use of a synteny-species reference genome). The previously identified scaffold containing the inversion polymorphism (scaffold28/NW_015090842.1 [6]) is a clear outlier.

#### 3.5 Output plots type 4 from analyses with fewer samples

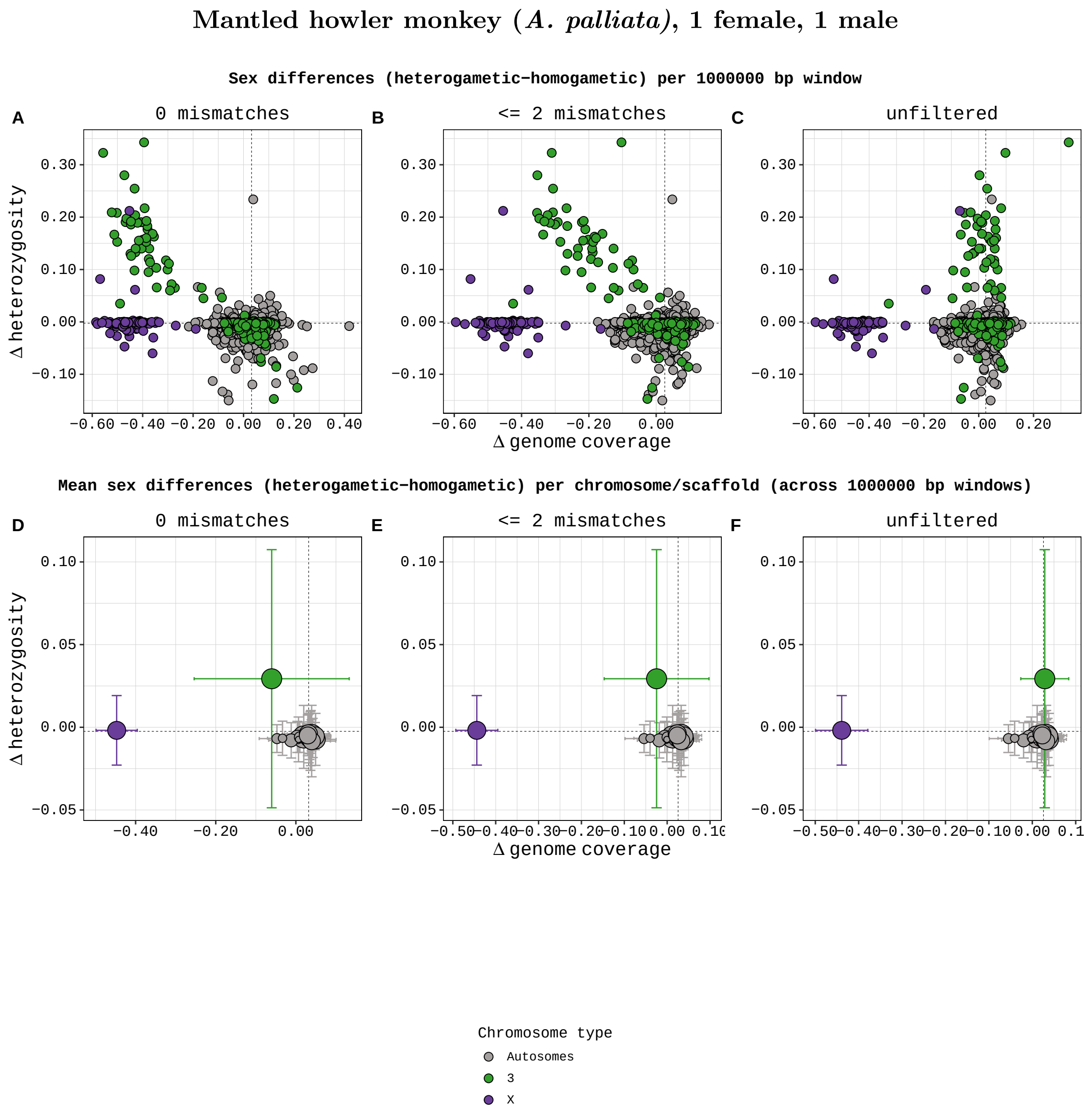


Supplementary Figure 14: **A-C)** Sex differences in genome coverage and heterozygosity for all 1 Mb genome windows. **D-F)** Mean (± SD) sex differences in genome coverage and heterozygosity per chromosome/scaffold, calculated from the 1 Mb genome windows. Dashed lines mark the genome-wide median across all 1 Mb windows. Data from 1 male and 1 female mantled howler monkey (*A. palliata*), analysed using findZX-synteny and human (*H. sapiens*) as a synteny-species reference genome. The previously identified sex chromosomes in this species (X and 3 [4]) are clear outliers.

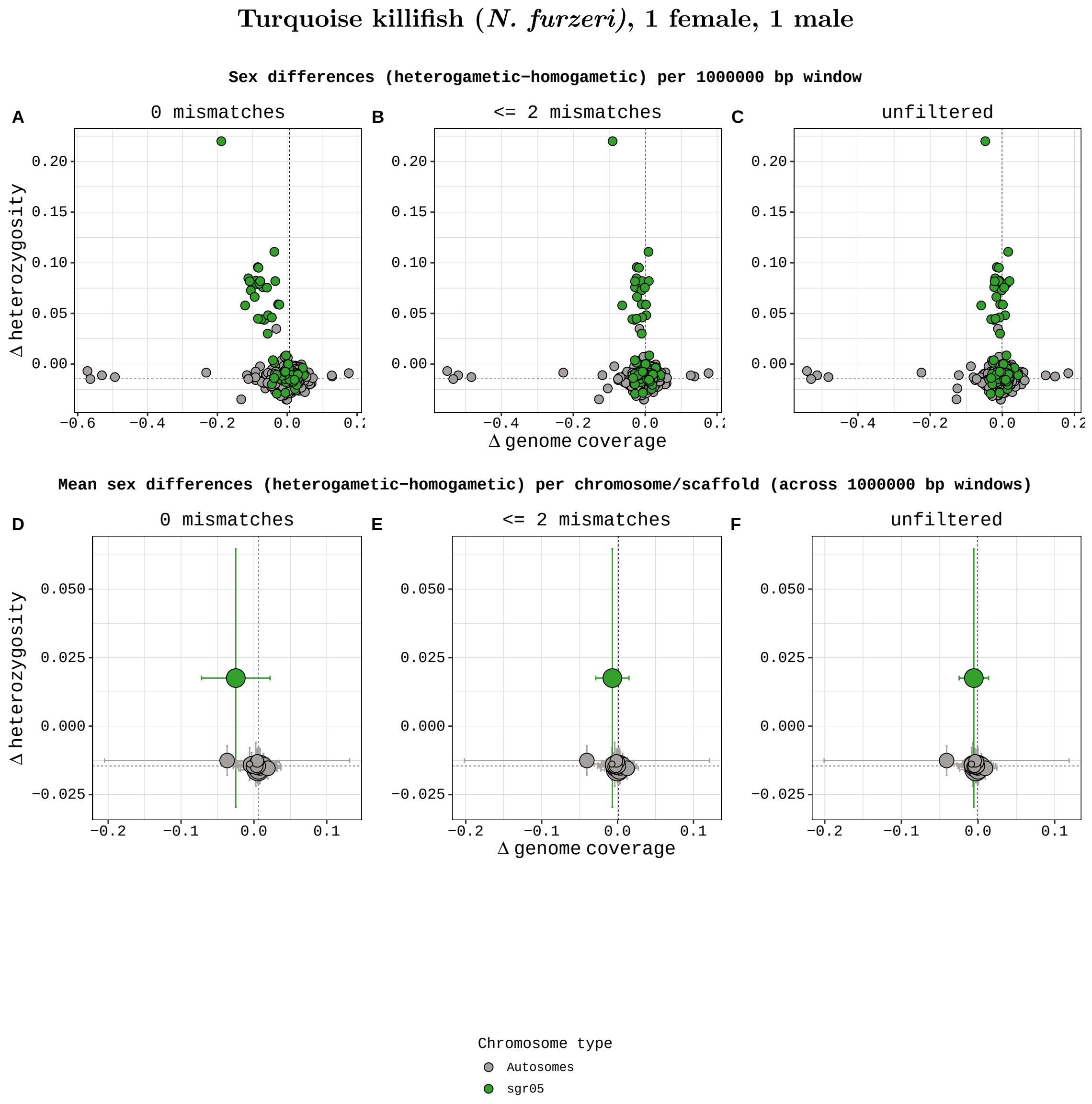


Supplementary Figure 15: **A-C)** Sex differences in genome coverage and heterozygosity for all 1 Mb genome windows. **D-F)** Mean (± SD) sex differences in genome coverage and heterozygosity per chromosome/scaffold, calculated from the 1 Mb genome windows. Dashed lines mark the genome-wide median across all 1 Mb windows. Data from 1 male and 1 female turquoise killifish (*N. furzeri*), analysed using findZX (without a synteny-species reference genome). The previously identified sex chromosome in this species (sgr05 [8]) is a clear outlier.

#### Guppy (*P. reticulata*), 1 female, 1 male

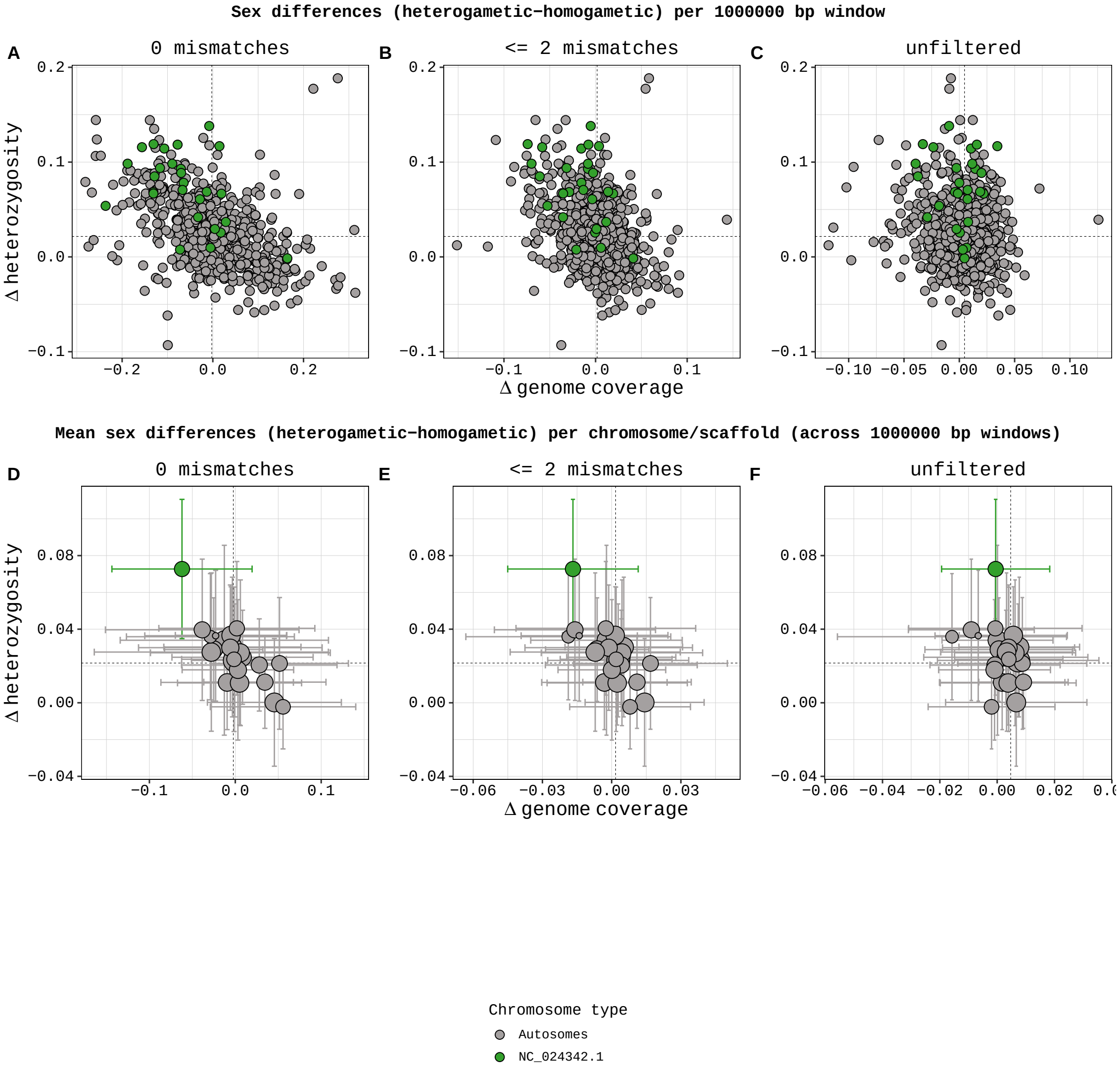


Supplementary Figure 16: **A-C)** Sex differences in genome coverage and heterozygosity for all 1 Mb genome windows. **D-F)** Mean (± SD) sex differences in genome coverage and heterozygosity per chromosome/scaffold, calculated from the 1 Mb genome windows. Dashed lines mark the genome-wide median across all 1 Mb windows. Data from 1 male and 1 female guppy (*P. reticulata*), analysed using findZX (without a synteny-species reference genome). The previously identified sex chromosome in this species (Chromosome 12/NC_024342.1 [10, 11]) is a clear outlier.

#### 3.6 Output plots type 4 from analyses of subsampled files

##### Mantled howler monkey (*A. palliata*), subsampled to 50% of the smallest sample

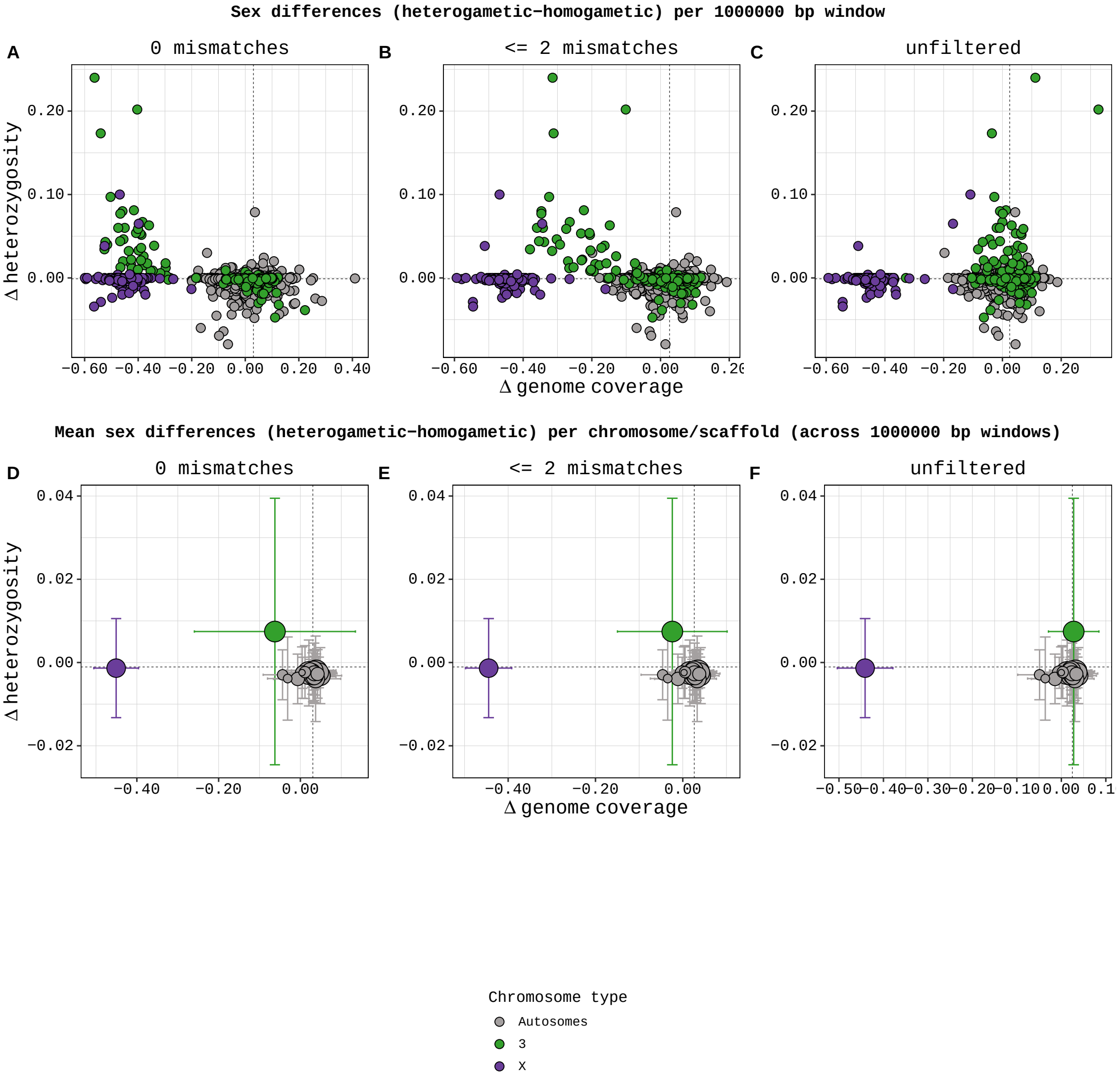


Supplementary Figure 17: **A-C)** Sex differences in genome coverage and heterozygosity for all 1 Mb genome windows. **D-F)** Mean (± SD) sex differences in genome coverage and heterozygosity per chromosome/scaffold, calculated from the 1 Mb genome windows. Dashed lines mark the genome-wide median across all 1 Mb windows. Data from 1 male and 1 female mantled howler monkeys (*A. palliata*), subsampled to 50% of the number of base pairs in the smallest of the fastq files. Analysed using findZX-synteny and human (*H. sapiens*) as a synteny-species reference genome. The previously identified sex chromosomes in this species (X and 3 [4]) are clear outliers.

#### 3.7 Output plots type 4 (and 1) from analyses using distantly related species as synteny-species

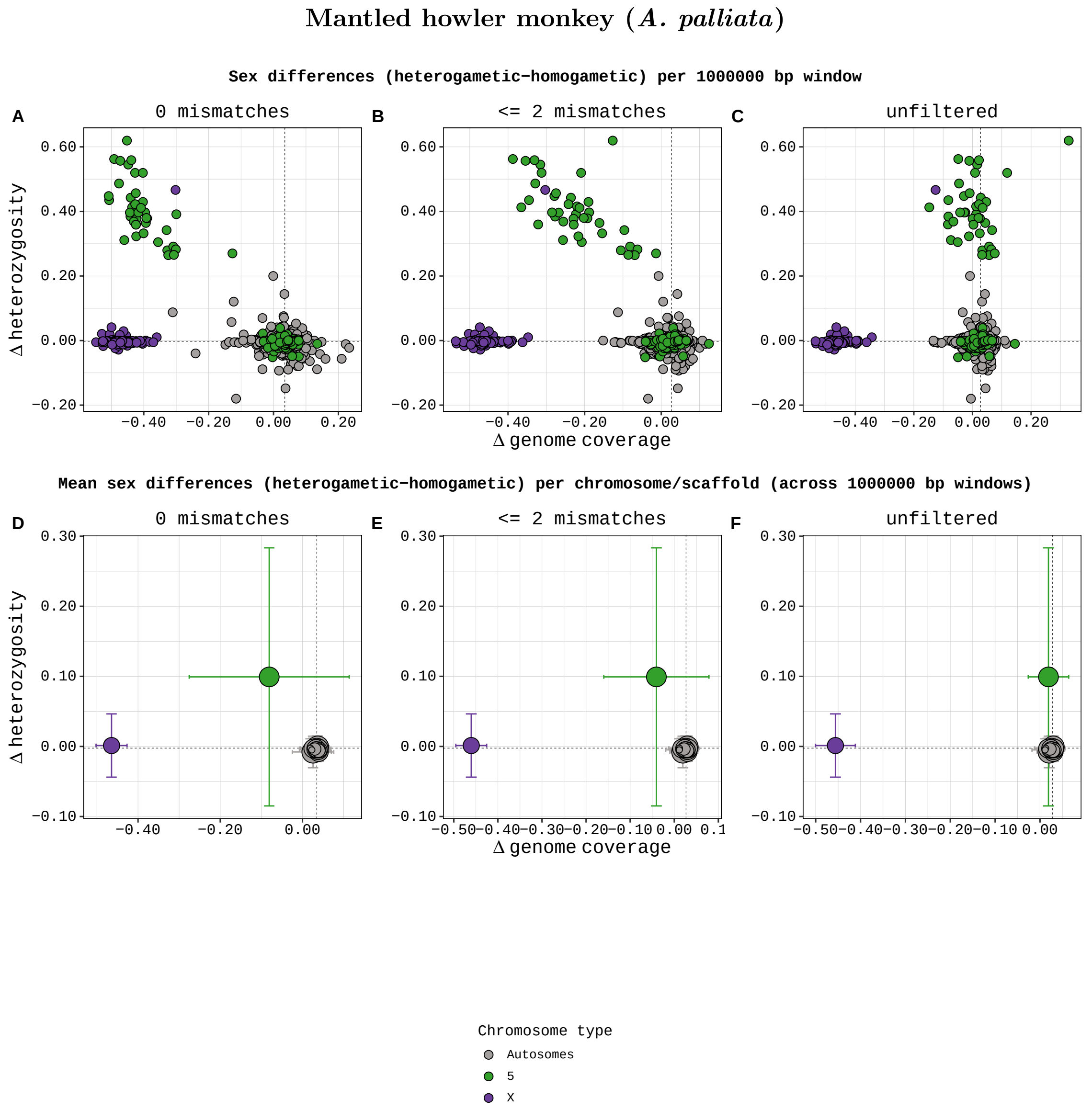


Supplementary Figure 18: **A-C)** Sex differences in genome coverage and heterozygosity for all 1 Mb genome windows. **D-F)** Mean (± SD) sex differences in genome coverage and heterozygosity per chromosome/scaffold, calculated from the 1 Mb genome windows. Dashed lines mark the genome-wide median across all 1 Mb windows. Data from 2 male and 2 female mantled howler monkeys (*A. palliata*), analysed using findZX-synteny and meerkat (*S. suricatta*) as a synteny-species reference genome. The previously identified sex chromosomes in this species (X and 5 [4]) are clear outliers.

#### Eurasian skylark (*A. alauda*)

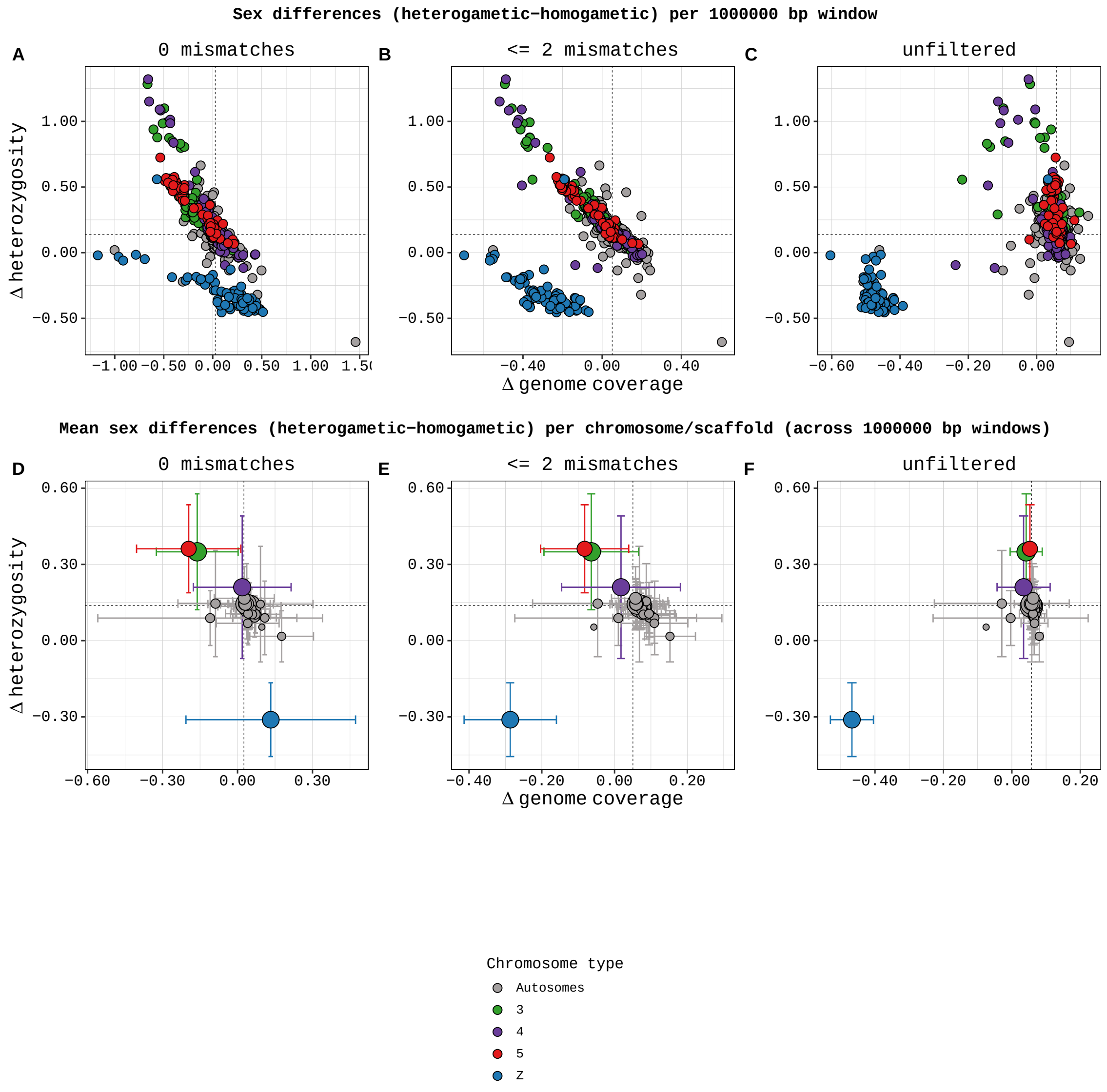


Supplementary Figure 19: **A-C)** Sex differences in genome coverage and heterozygosity for all 1 Mb genome windows. **D-F)** Mean (± SD) sex differences in genome coverage and heterozygosity per chromosome/scaffold, calculated from the 1 Mb genome windows. Dashed lines mark the genome-wide median across all 1 Mb windows. Data from 1 male and 1 female Eurasian skylark (*A. arvensis*), analysed with findZX-synteny and chicken (*G. gallus*) as a synteny-species. Three of the four previously identified sex chromosomes in this species are clear outliers (Z, 3 and 5 [1, 2]). Chromosome 4 is not a clear outlier, as a result of the sex-linked part of this chromosome only constituting 10 Mb of this chromosome (91 Mb in total). Supplementary Figure 20 show the same data as in this plot, but across chromosome positions in the chicken genome. In this plot, the beginning of chromosome 4 (which is homologous to chromosome 4A in the zebra finch) show a sex-linked pattern. The analysis was performed using a "consensus reference genome".

#### Eurasian skylark (*A. alauda)*

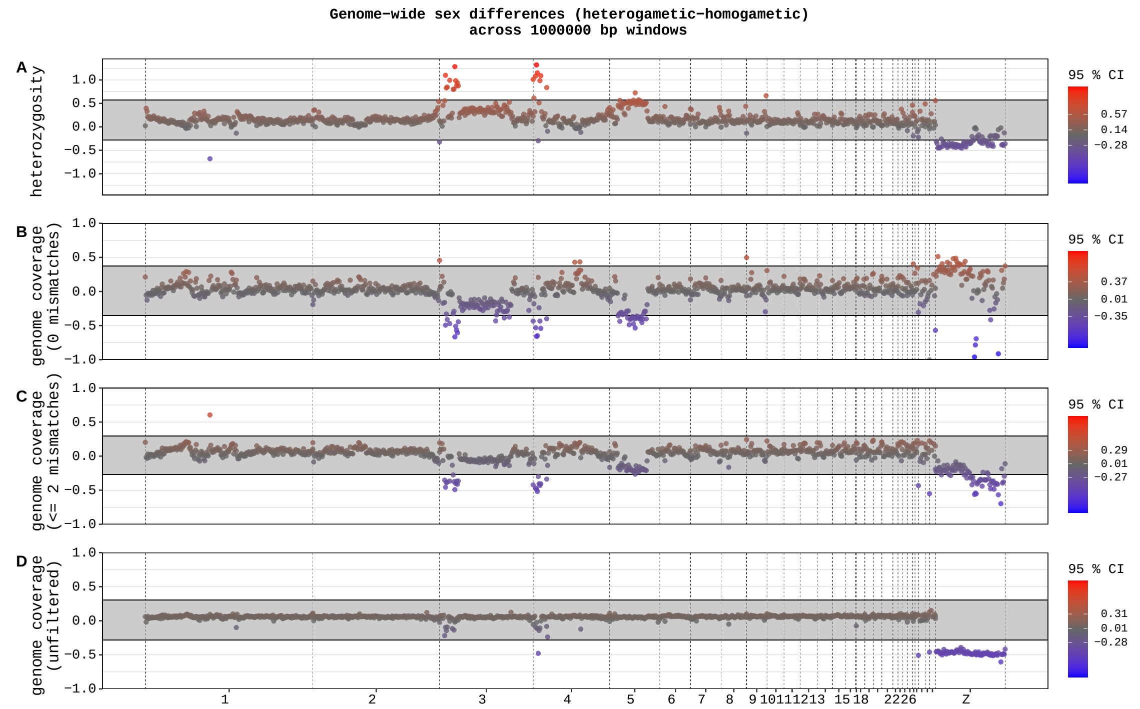


Supplementary Figure 20: Sex differences in genome coverage and heterozygosity values (1 Mb windows) for Eurasian skylark, plotted along chromosome positions in the chicken genome. The four rows show: (**A**) heterozygosity, and genome coverage with (**B**) strict filtering (0 mismatches allowed), (**C**) intermediate filtering ($\leq$2 mismatches) and (**D**) no filtering of mapped reads (unfiltered). The previously identified sex chromosomes in this species (Z, 3, 4 and 5 [1, 2]) are clear outliers. The analysis was performed using a "consensus reference genome".

#### Ruff (*C. pugnax*)

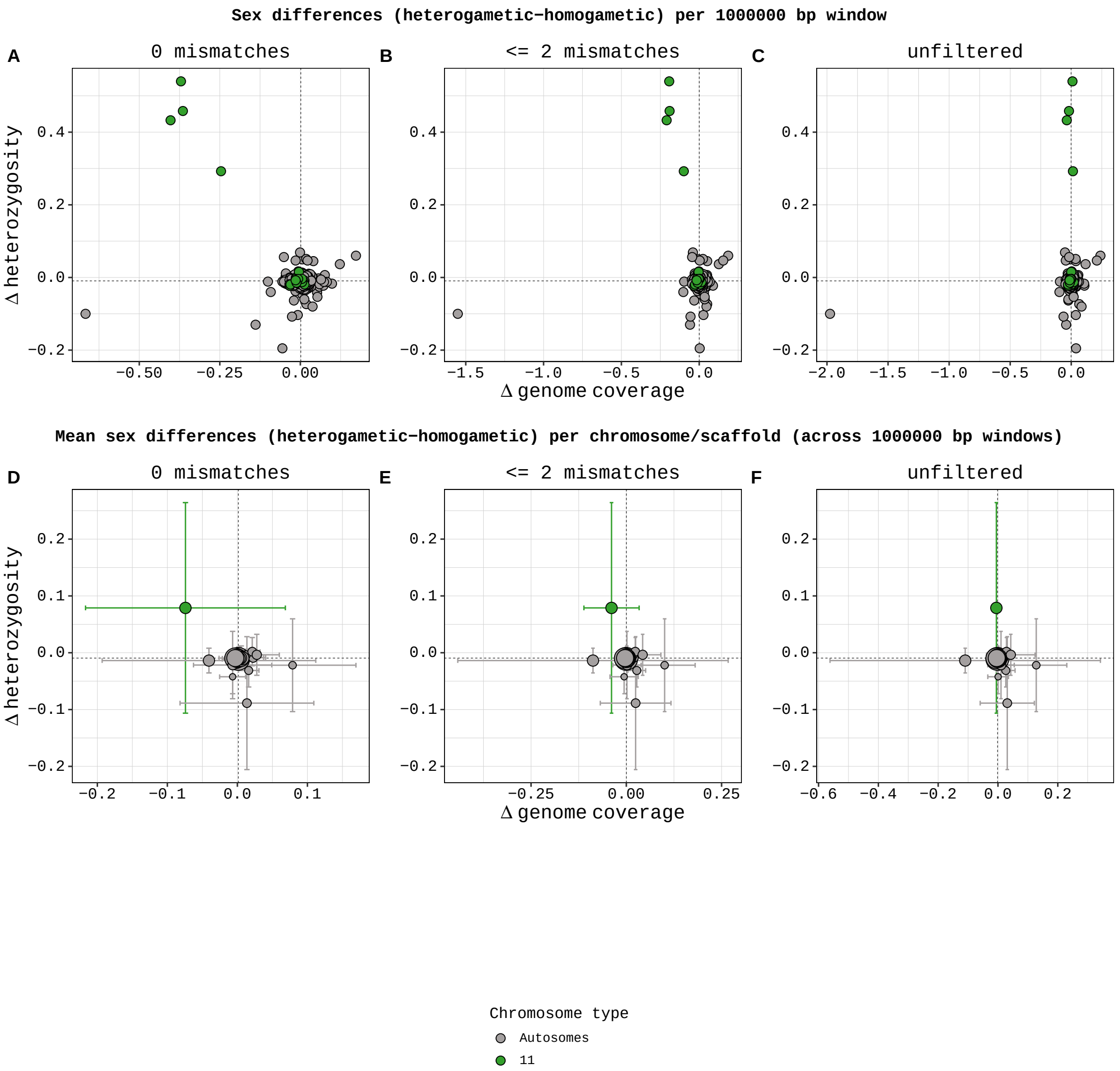


Supplementary Figure 21: **A-C)** Morph differences in genome coverage and heterozygosity for all 1 Mb genome windows. **D-F)** Mean (± SD) morph differences in genome coverage and heterozygosity per chromosome/scaffold, calculated from the 1 Mb genome windows. Dashed lines mark the genome-wide median across all 1 Mb windows. Data from 3 male ruffs (*C. pugnax*); 2 faeder males (heterozygotic inversion) and 1 resident male (homozygotic inversion).The data was analysed using findZX-synteny with chicken (*G. gallus*) as a synteny-species reference genome. Chromosome 11, which contain the inversion polymorphism [6], is an outlier.

### Turquoise killifish (*N. furzeri*)

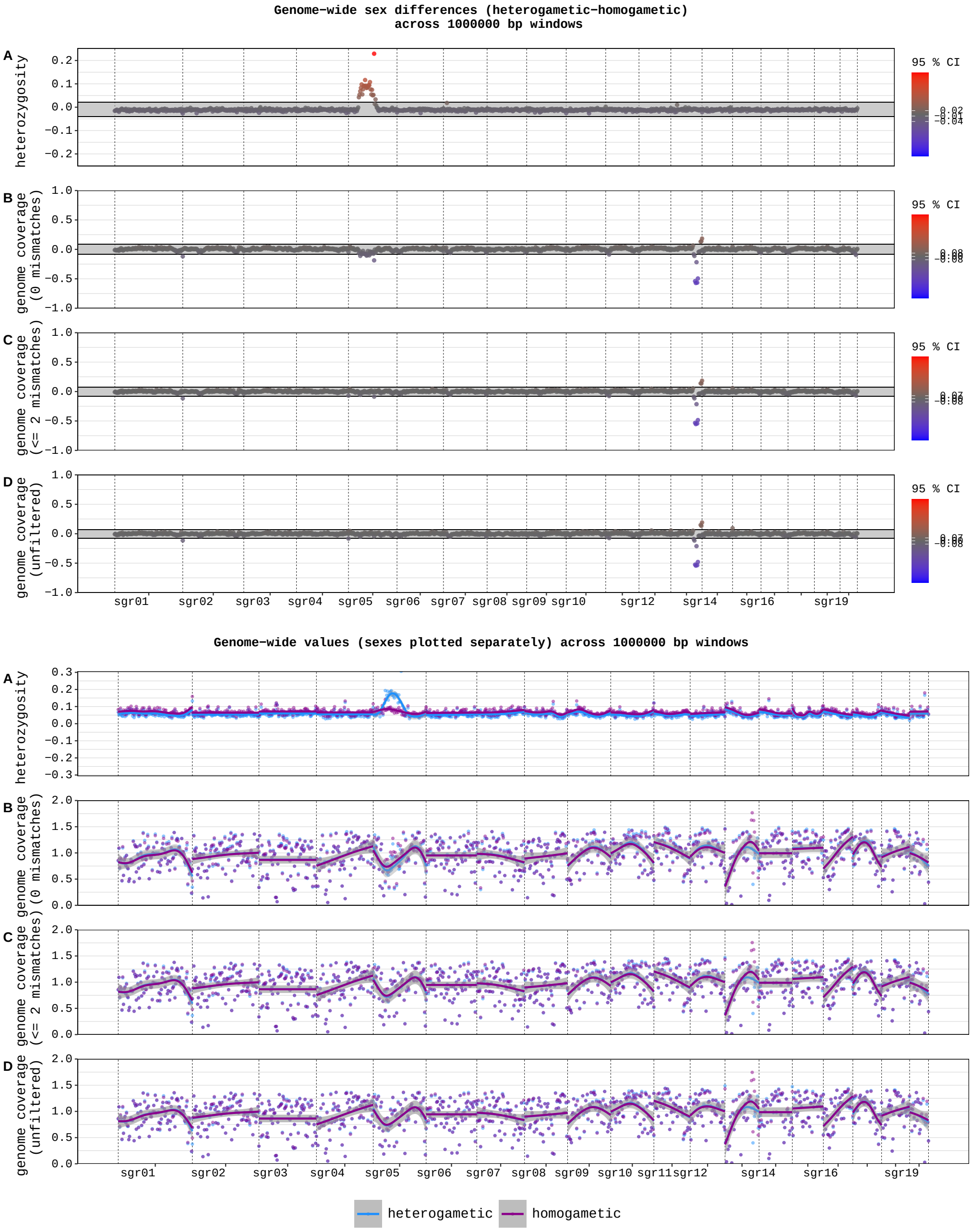


Supplementary Figure 22: Genome-wide heterozygosity and genome coverage values from 2 male and 1 female turquoise killifish (*N. furzeri*), analysed using findZX. In the top figure, sgr13 shows a reduction in genome coverage in the heterogametic sex compared to the homogametic one. In the bottom figure, however, there is no reduction in genome coverage in the heterogametic sex for sgr13 compared to the genome average. The bottom figure shows that the homogametic sex has slightly elevated genome coverage values in this region compared to the rest of the genome, which likely produced the pattern we see in the top figure. When interpreting results provided by this pipeline, it is important to look at all the plots in order to avoid misinterpretations (see Discussion).

**Phylopic/Wikimedia commons credits:**
The guppy silhouette (Figure 5g) was created by Josefine Bohr Brask and shared via phylopic.org under a CC By 3.0 license (https://creativecommons.org/licenses/by/3.0/). Not modified from its original state.

The ruff silhouette (Figure 5i) was created by Alexandre Vong and shared via phylopic.org under a CC By 3.0 license (https://creativecommons.org/licenses/by/3.0/). Not modified from its original state.

Silhouettes in Figure 5a-f,h were downloaded from phylopic.org and were without copyright restrictions.
